# Genotype Imputation and Reference Panel: A Systematic Evaluation

**DOI:** 10.1101/642546

**Authors:** Wei-Yang Bai, Xiao-Wei Zhu, Pei-Kuan Cong, Xue-Jun Zhang, J Brent Richards, Hou-Feng Zheng

## Abstract

Here, 622 imputations were conducted with 394 customized reference panels for Han Chinese and European populations. Besides validating the fact that the imputation accuracy could always benefit from the increased panel size when the reference panel was population-specific, the results brought two new thoughts as follows. First, when the haplotype size of reference panel was fixed, the imputation accuracy of common and low-frequency variants (MAF>0.5%) decreased while the population-diversity of reference panel increased, but for rare variants (MAF<0.5%), a fraction of diversity (<20%) of panel could improve the imputation accuracy. Second, when the haplotype size of reference panel was increased with extra population-diverse samples, the imputation accuracy of common variants (MAF>5%) for European population could always benefit from the expanding sample size. But for Han Chinese population, the accuracy of all imputed variants reached the highest when reference panel contained a fraction of extra diverse sample (15%∼21%). In addition, we evaluated the existing reference panels such as the HRC and 1000G Phase3 and CONVERGE. For European population, HRC was the best reference panel. For Han Chinese population, we proposed an optimum constituent ratio for the Han Chinese imputation if researchers would like to customize their own sequenced reference panel, but a high quality and large-scale Chinese reference panel was still needed. Our findings could be generalized to the other populations with conservative genome, a tool was provided to investigate other populations of interest (https://github.com/Abyss-bai/reference-panel-reconstruction).

**Highlights (Key points):**

1. A total of 394 reference panels were designed and customized by three strategies, and large-scale genotype imputations were performed with these panels for systematic evaluation in Han Chinese and European populations.
2. The accuracy of imputed variants reached the highest when reference panel contains a fraction of extra diverse sample (15%∼21%) for Han Chinese population, if the haplotype size of the reference panel was increased with extra samples, which is the most common cases.
3. The imputation accuracy showed the different trends between Han Chinese and European populations. In a sense, the European genome may more diverse than Han Chinese genome by itself.
4. Existing reference panels were not the best choice for Chinese imputation, a high quality and large-scale Chinese reference panel was still needed.

## Introduction

As a cost-efficient way of genotyping variants, imputation has become a standard approach in genome-wide association studies (GWAS) in the past decade. It is achieved by using known haplotypes in a population to infer initially-untyped genetic variants for testing association with a trait of interest[1], thereby allowing to overcome one major limitation of SNP genotyping arrays. In generally, SNP array only contains a small fraction of human genetic variation (10^5^–10^6^), genotype imputation makes these low-density genetic variants array become a higher one (10^7^– 10^8^). A higher-resolution view of a genetic region can provide many advantages for population genetics research, such as guiding fine-mapping by increasing the chances of identifying a causal variant[2], facilitating the combination of results across studies in meta-analysis[3, 4], and increasing the power to detect an association signal[5, 6]. Since genotype imputation carries such potentials, accuracy of imputed-variant is crucial. Many studies have illustrated the accuracy and reliability of genotype imputation in common variants (MAF > 5%)[7–9]. Compared to common variants, rare variants are often population specific and tend to have low levels of pairwise linkage disequilibrium with other sites, but more likely to be associated with dramatic functional consequences[10, 11]. More and more rare and low-frequency variants were discovered to be associated with serious diseases[3, 12–14]. However, keep the imputation accuracy of rare and low-frequency variants at a reliable level is still a challenge[15].

Several imputation tools have been developed during the last decade, most of them employ the hidden Markov model (HMM) as their engine, such as IMPUTE, MaCH and Minimac series[16–18]. Although the algorithms of imputation tools are constantly updated, the main purpose of them is to reduce the compute pressure of server, the assistance they can provide in improving accuracy of imputation of rare variants is very limited. Imputation reference panel as the haplotype patterns and information carrier for inference of untyped genetic variants, its composition and size are far more crucial influential factors for imputation accuracy[19].

Since the International HapMap 3 Project was accomplished in 2010[20], more and more whole-genome sequencing (WGS) data were produced for public use. The quality of genotype imputation has benefited from the increase of genetic information in these publicly available reference panels data[21, 22], one of the most famous and widely used reference panel is from the 1000 Genomes Project (1000G)[23]. The 1000G Project Phase 3 identified more than 84.4 million single nucleotide polymorphisms (SNP) from 2,504 individuals which collected from 26 worldwide populations, each population contains 61∼113 individuals. All of the 26 populations were divided into 5 groups (AFR, African; AMR, Ad Mixed American; EAS, East Asian; EUR, European; SAS, South Asian). Besides the 1000G Project, there are some more population-specific reference panels. Examples include the UK10K Project[24] (3781 British sequenced at 7× depth of coverage) and the Genome of the Netherlands[25] (GoNL, 250 Dutch parent–offspring families sequenced at 14× depth). Recently, a large combined haplotype reference panel named the Haplotype Reference Consortium (HRC) was formed, it consists of 64,976 haplotypes at 39,235,157 SNPs with the minor allele count (MAC) greater or equal to 5, and it will collect more WGS data in future[26]. In 2017, 11,670 genomes of Chinese sequencing project called The China, Oxford and Virginia Commonwealth University Experimental Research on Genetic Epidemiology (CONVERGE) was published, but only ∼22 million high quality SNPs were identified because of its low-coverage sequencing depth (1.7×)[27].

There are many factors that affect imputation accuracy of rare variants, such as density of genotyping array, ancestry diversity of GWAS data, as well as sequencing depth, haplotype size and diversity of reference panel[1]. In general, for genetically diverse populations such as Hispanics/Latinos in the USA, a corresponding diverse reference panel will improve the imputation accuracy[28]. For an ancestry-specific GWAS data, such as Southeast Asian[29] and African ancestry[8], using the corresponding specific reference would gain more accuracy because of the same genetic background. However, another study found that the accuracy of imputation of low-frequency variants can benefit from the reference diversity, independent of reference haplotype size[30]. It is widely accepted that the haplotype size is a key factor in a particular reference panel, but mostly, expanding the reference panel size means to introduce more population diversity.

Therefore, it is remaining unclear that the relationship between imputation accuracy of rare variants and composition of reference panels and how to maximize the imputation accuracy. Here, we proposed a much rigorous and systematic design to evaluate the relationships between imputation performance and haplotype diversity and size of reference panel for Han Chinese and European populations, by using 394 customized reference panels and by performing 622 imputations. Besides, we evaluated the rare variants imputation performance of HRC, 1000G and CONVERGE reference panels for both Han Chinese and European populations

## Methods

### Sample datasets and genotyping

In this study, we used Han Chinese and European samples as GWAS sets. All Han Chinese samples, which consist of 2,360 individuals, were obtained from multiple regions in central and southern China[31]. The Illumina Human610-Quad (610K) BeadChip was employed for genotyping analysis based on the Genome Reference Consortium Human build 36 (GRCh36) and a total of 598,821 SNPs were identified. The European dataset was obtained from TwinsUK Project (http://www.twinsuk.ac.uk/). All 3,461 European individuals were genotyped by using the 610K BeadChip[32] which was the same as the Han Chinese genotyping array, and the genome annotation was based on GRCh36.

### Quality control and pre-phasing

We first updated the genome assembly version of genotyping array variants from build 36 to 37. For all sample datasets, we performed a stringent quality control (QC) with four steps. Step one, we retained autosomal bi-allelic SNPs with missing call rates <= 5% and samples with missing call rates <= 2% of data by using PLINK v1.9[33]. Step two, the pairwise genetic relationship matrix between all samples was calculated by GCTA v1.91[34] using common variants with MAF > 10%, and individuals with pairwise genetic relationship coefficient > 0.025 will be thought to be cryptically related. We then randomly selected 2000 unrelated individuals for both Han Chinese and European samples sets. Step three, we downloaded the legend file of 1000G Phase 3, and used the EAS (East Asians) and EUR (Europeans) populations as the reference to check our Han Chinese and European data respectively by a Perl scripts of checking tools (www.well.ox.ac.uk/~wrayner/tools/). We checked if any SNP ID or genome position was mismatched with reference panel, if yes, the SNP was removed, and we corrected the allele switch and strand flip in the GWAS sets, and we removed SNPs whose allele frequency difference with reference was larger than 0.2. Last step, we excluded the SNPs with missing call rates > 5% again, and excluded those deviating from Hardy–Weinberg equilibrium (HWE) at P < 1 × 10^−6^. In order to study the imputation accuracy of very rare variants, we retained all SNPs in Chinese sample dataset. And note that SNPs in European were all with MAF > 0.5%. Finally, 516,410 overlapped variants between Han Chinese and European in total were used as the study data.

To reduce the computed pressure of the subsequent large-scale genotype imputations, we phased Han Chinese and European datasets by using SHAPEIT v2.9[35] with the default settings in local server. And we checked our QCed data in Michigan imputation server[16].

### Reference panels re-construction by haplotype size

Twenty-four out of the 26 populations in the 1000G Phase 3 reference panel with sample size greater than 85 were selected to study the relationship between haplotype size and imputation accuracy, that 3 were from AMR (Ad Mixed American), 6 were from AFR (African) and other 15 were from EAS (East Asian), SAS (South Asian) and EUR (European) respectively. We randomly and successively extracted 25, 50, 65, and 85 samples from each population alone (Figure 1a) and customized them into 4 size gradients, which were 50, 90, 130 and 170 haplotypes, and the higher gradient sets contained all haplotypes of the lower one. The bcftools was employed here (https://samtools.github.io/bcftools/). These customized 96 (24 times 4) panels were then used for imputation of Han Chinese and European sample sets by Minimac3 in local server, respectively (Figure 1a).

**Figure 1.**
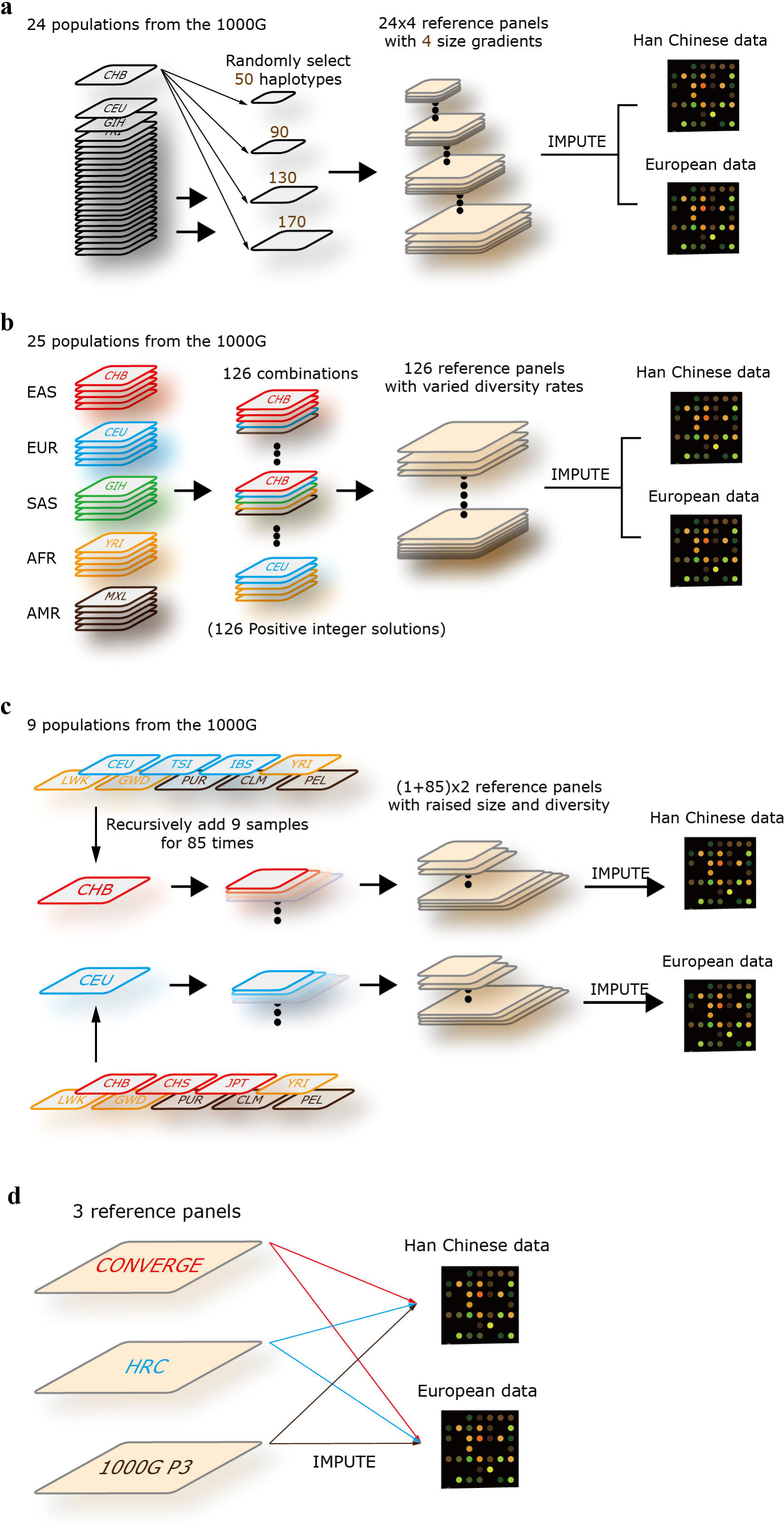
Research design. **(a)** The design of imputation accuracy vs. reference panel size. 24 worldwide population of the 1000G Phase3 were selected (sample size > 85), such CHB, CEU and GIH. For each population, the haplotypes were extracted and customized as reference panels with 4 sizes gradients (50, 90, 130, 170 haplotypes). Totally, 96 reference panels (24 times 4) were formed, and then the imputation were performed for the Han Chinese and European chip data in local server. **(b)** The design of imputation accuracy vs. reference panel diversity. In this part, the size of reference panels was fixed to 640 haplotypes. 25 populations of the 1000G Phase3 were selected and categorized into 5 groups (EAS, EUR, SAS, AFR and AMR, see Methods), each group included 5 populations and each population contained 64 samples. The 5 groups of population that we considered corresponded to a set of vectors (*i_1_ to i_5_*), and solved the function of *i_1_* + *i_2_* + *i_3_* + *i_4_* + *i_5_* = 5. We got 126 positive integer solutions in total, which represented 126 combinations. Finally, 126 reference panels were formed and the imputations were performed for the Han Chinese and European chip data in local server. **(c)** The design of imputation accuracy vs. reference panel size & diversity. 12 populations were selected. 9 of them were diverse to Han Chinese and European population respectively. Then, the CHB and CEU were used as basic panel, the diverse samples were recursively added to them, 9 samples per time and 85 times in total. Finally, two reference panels set were formed, and each group included 86 reference panels (1+85). The imputations were then performed for the Han Chinese and European chip data in local server. **(d)** Imputation for the Han Chinese and European population using 1000G, HRC and CONVERGE reference panels.

### Reference panels re-construction by population diversity

We categorized 5 groups corresponded with EAS, EUR, AFR, AMR and SAS of populations distribution in the 1000G Phase 3 to study the relationship between imputation accuracy and diversity of reference panel (Figure 1b). Each group consists of 5 populations in sequence and each population contains 64 samples (Supplementary Table 1). Note that AFR group has 7 populations in the 1000G, we excluded the ‘ASW’ (Americans of African Ancestry in SW USA) in AFR because of its small sample size, and re-categorized the ‘ACB’ (African Caribbeans in Barbados) into AMR group in this study (see Discussion). The 5 groups of population that we considered corresponded to a set of vectors (*i_1_* to *i_5_*), then we solved the function:

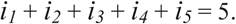

126 non-negative integer solutions were obtained in total. These solutions corresponded to 126 different combinations of 25 populations in 5 groups, the reference panels were then constructed based on these combinations. For these panels, we set six levels of population diversity based on the number of populations of EAS or EUR, the diversity degree was raised from level_0_ to level_5_, for example, the solution (*i_1_, i_2_, i_3_, i_4_, i_5_*) = (2, 3, 0, 0, 0) means a reference panel consists of the first 2 populations in EAS, and the first 3 populations in EUR (Supplementary Table 1), and no AFR, AMR or SAS populations was included. Therefore, this panel was diverse for EAS at level_3_ and for EUR at level_2_. Each panel contains 640 haplotypes. The imputations were performed for Han Chinese and European sample sets with 126 different diversity panels (Supplementary Table 2) by using Minimac3 in local.

### Reference panels re-construction by both haplotype size and population diversity

In this part, a series of reference panels were customized with haplotype size and population diversity constantly changed (Figure 1c). We took these two factors together as arguments to investigate the pattern of imputation accuracy variation. First, we extracted CHB (Chinese in Beijing) and CEU (Utah Residents with Northern and Western European Ancestry) population samples from 1000G Phase 3 according to the ancestry to our GWAS study sets. Besides, we also extracted other 10 populations included 3 AMR populations (PUR, CLM, PEL), 3 AFR populations (YRI, LWK, GWD), 2 EUR populations (TSI, IBD) and 2 EAS populations (CHS, JPT). The CHB and CEU population contains 103 and 99 samples respectively, and each of other 10 populations contains at least 85 samples. Then, we took CHB and CEU samples as two basic panels, and added other population samples to them constantly. To ensure that no individuals from corresponding specific group were involved in CHB-based and CEU-based panel, we used different adding strategies. For CHB-based panel, we chose the adding-populations from AMR, AFR and EUR groups. For CEU-based panel, we chose the adding-populations from AMR, AFR and EAS groups (Figure 1c), each group contained 3 populations, and then we respectively took one individual from these 9 populations per time, and recursively added them to basic panel for 85 times in total (Figure 1c). Finally, we got 172 imputation reference panels, half of them were CHB-based and another half were CEU-based. These panels were then used for Han Chinese and European sample sets imputation by using Minimac3 in local server, respectively.

### HRC, 1000G and CONVERGE reference panels

The HRC was the largest reference panel for genotype imputation currently and mainly consist of European population samples[26]. The 1000G sample was consist of 26 worldwide populations and made it the most diverse reference panel[23]. The CONVERGE (The China, Oxford and Virginia Commonwealth University Experimental Research on Genetic Epidemiology) was a Chinese-specific panel with 11,670 genomes with low depth (1.7×)[27]. These three reference panels were assessed in our study (Figure 1d). Minimac3 was employed for genotype imputation in our study because of its advanced performance[16]. We converted all reference sets from common VCF format into Minimac3 specialized M3VCF format which require lesser space and are faster to read than VCF file while importing data.

A basic statistics of variants was performed first between HRC, 1000G and CONVERGE reference panels. We used remote and local ways to impute our data, since the complete haplotypes set of the HRC was not available for downloading yet, and the CONVERGE-based imputation was performed using Minimac3 in local server. For consistency of imputation tools, the Michigan Imputation Server was employed for remote HRC-based imputation. The 1000G-based imputation was performed in both ways (Michigan Imputation Server and local imputation server) to test the comparability of results of two different servers. Note that the 1000G panel had two versions, one included singletons and another one not, the latter was mainly used here (see Discussion). All imputations were conducted with default settings.

### Evaluation of imputation accuracy

In this study, we employed Minimac3 statistics including R^2^ and empirical-R^2^ (EmpR^2^) to evaluate genotype imputation quality and accuracy. Both R^2^ and EmpR^2^ value of each sites can be obtained from imputation results. R^2^ was the estimated value of the squared correlation between imputed genotypes and true, unobserved genotypes. Since true genotypes were not available, this calculation was based on the idea that poorly imputed genotype counts will shrink towards their expectations based on population allele frequencies alone[16]. R^2^ was defined as:

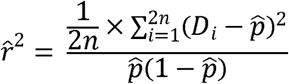

Where *p* was the alternate allele frequency and *D_i_* was the imputed alternate allele probability at the *ith* haplotype and *n* was the number of GWAS study samples.

For each site that were genotyped in the study samples, Minimac3 can calculates a special imputed dosage by hiding all known genotypes for that site. This imputed value is called leave-one-out dosage (LooDosage) and was used to calculate EmpR^2^ by directly calculating the Pearson correlation coefficient between LooDosage and known genotypes. Compared to R^2^, EmpR^2^ was more powerful and effective to evaluate imputation accuracy, and can only be calculated in genotyped sites. In our study, we set a strict threshold for ‘well-imputed’ sites, which the Minimac3 R^2^ (imputation quality) had to reach at least 0.8, and used Minimac3 EmpR^2^ to measure the imputation accuracy.

## Results

### Imputation accuracy versus haplotype size of panel

We conducted a strict QC for Han Chinese and European genotyping array data sets in local, and checked the data after QC by using Michigan imputation server. The allele-frequency showed a strong correlation between GWAS sets and EAS or EUR data from the 1000G Phase 3 reference panels, which r^2^=0.991 for Han Chinese GWAS data set and r^2^=0.994 for European GWAS data set, respectively (Supplementary Figure 1). After QC, 516,410 overlapped sites in 2,000 unrelated individuals retained respectively for both populations.

It is generally accepted that genotype imputation accuracy can benefit from increasing panel size. In this study, we validated this point in a more systematic approach. We performed 192 imputations for Han Chinese and European GWAS data in this part. 24 worldwide populations from 1000G Phase 3 were customized as reference panel, each population was transformed into 4 gradients according to the number of haplotypes. All different population panels showed the consistent results that imputation accuracy increased with panel’s haplotypes size for both Chinese and European datasets (Figure 2a and 2b). For Han Chinese and European samples, the average accuracy of all genotyped-variants reached the highest when we used CHB and CEU population samples as the reference panel, respectively. To obtain the more distinct comparison between gradientized reference panels, we divided variants into five MAF bins including: 5% <= MAF < 100%, 1% <= MAF < 5%, 0.5% <= MAF < 1%, 0.1% <= MAF < 0.5% and 0.025% < MAF < 0.1%. The imputation accuracy in different MAF bins all showed an increasing trend when haplotypes size constantly augmented (Figure 2c and 2d). Besides, we counted well-imputed (R^2^ > 0.8) number of variants, the results showed that well-imputed variants number also raised with haplotypes size (Supplementary Figure 2a and 2b). These results validated the fact that the imputation quality could be improved by the haplotype size of reference panel.

**Figure 2.**
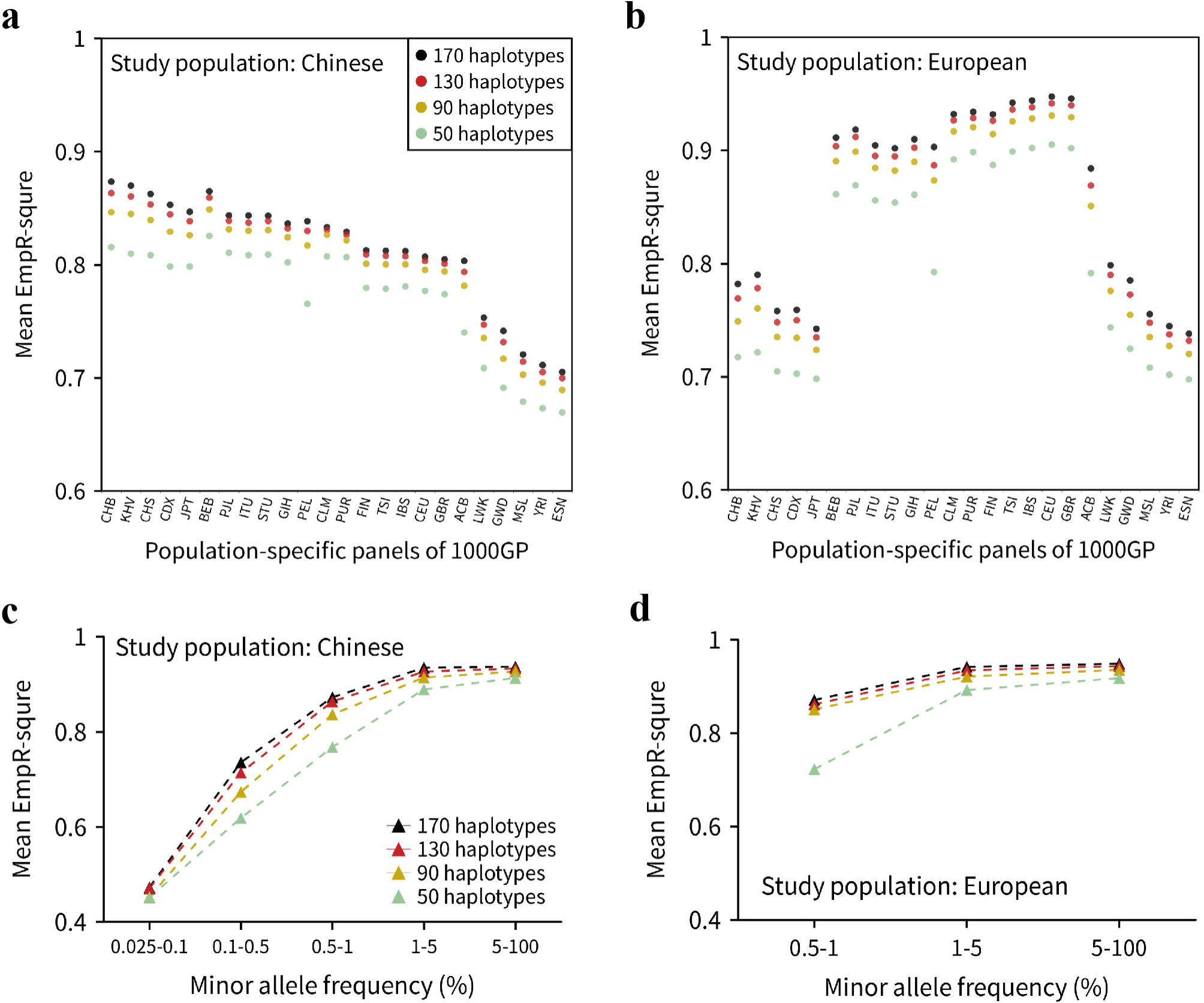
Imputation accuracy for the reference panels with four haplotype size gradients. Overall imputation accuracy for **(a)** the Han Chinese and **(b)** European using 24 worldwide populations of the 1000G Phase3 as reference panels. For each reference panel, the average EmpR^2^ (measuring the imputation accuracy) was plotted with four haplotype sizes (50, 90, 130 and 170). **(c)** Imputation accuracy for the Han Chinese in 5 different MAF bins using CHB (Han Chinese ancestry) population as the reference panel. **(d)** Imputation accuracy for the European using CEU (European ancestry) population as the reference panel. Only three MAF bins of EmpR2 of variants were plotted since the (upper) was for the Han Chinese imputation and the asterisk marked group (lower) was for the European imputation.

The average EmpR^2^ results of comparing the Han Chinese imputation and European imputation showed that the European population could be more accurately imputed when the corresponding population reference panel was used (increasing from 0.82 to 0.87 for Han GWAS and CHB, and from 0.91 to 0.95 for European GWAS and CEU while haplotype size increased from 50 to 170). The same pattern was also showed in heat map, that European GWAS imputation using reference panels from EUR were much redder than Han GWAS imputation using reference panels from EAS (Figure 3).

### Imputation accuracy versus population diversity of panel

In this part, we constructed 126 “diversity” reference panels using the five population groups of the 1000G (Supplementary Table 2), and performed 252 imputations in total. The size of each reference panel was *fixed* to 640 haplotypes. The overall average EmpR^2^ decreased from 0.92 to 0.84 for Chinese samples and from 0.96 to 0.93 for European samples while diversity changed from minimum degree to maximum degree (level_0_ to level_5_). We further divided imputed variants into 5 MAF bins, as shown in Figure 4a and 4b, the imputation accuracy of variants with MAF >= 0.5% showed the decreasing trend when diversity degree raised. However, for the rare variant imputation (MAF < 0.5%) in Han Chinese population, the accuracy increased when the diversity degree raised from level_0_ to level_1_. These results suggested that, when the haplotype size was fixed, the more that a reference panel specific to the study population, the more accurately that it could impute for common variants and low-frequency variants. But for rare variants (MAF < 0.5%), a little diverse fraction of population of reference panel could benefit the imputation accuracy.

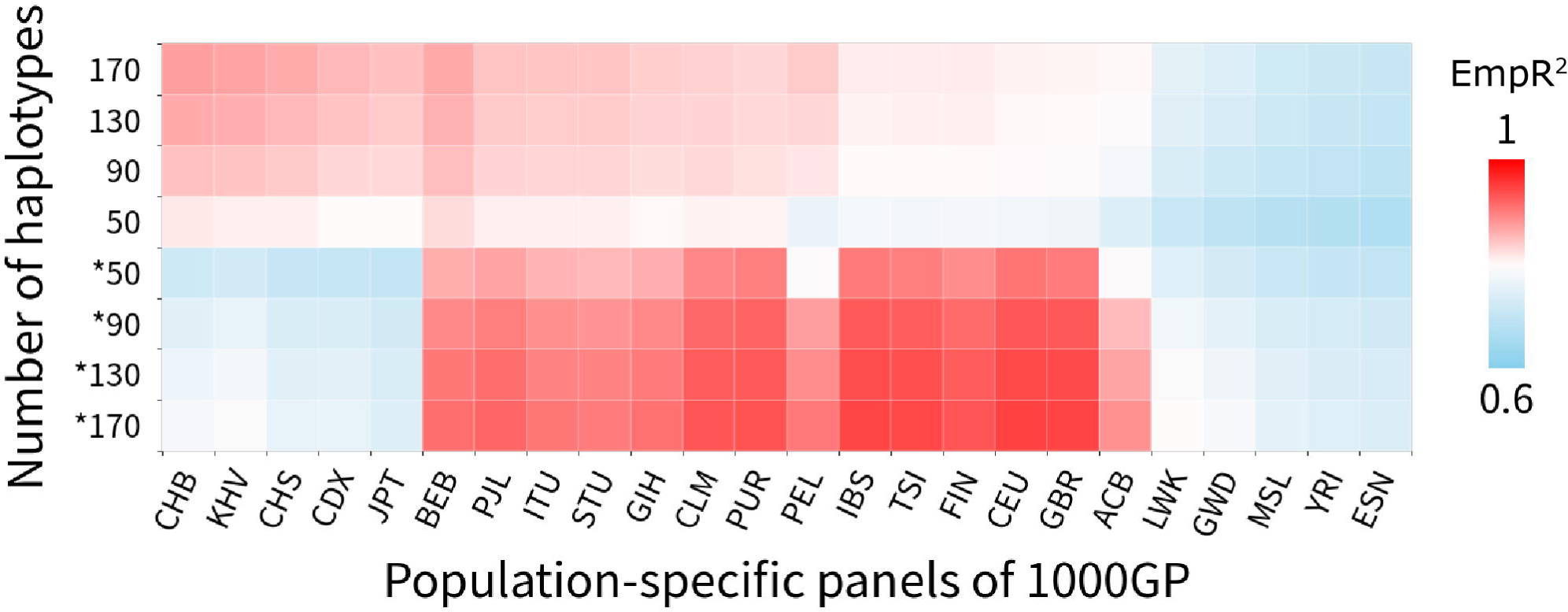

**Figure 4.**
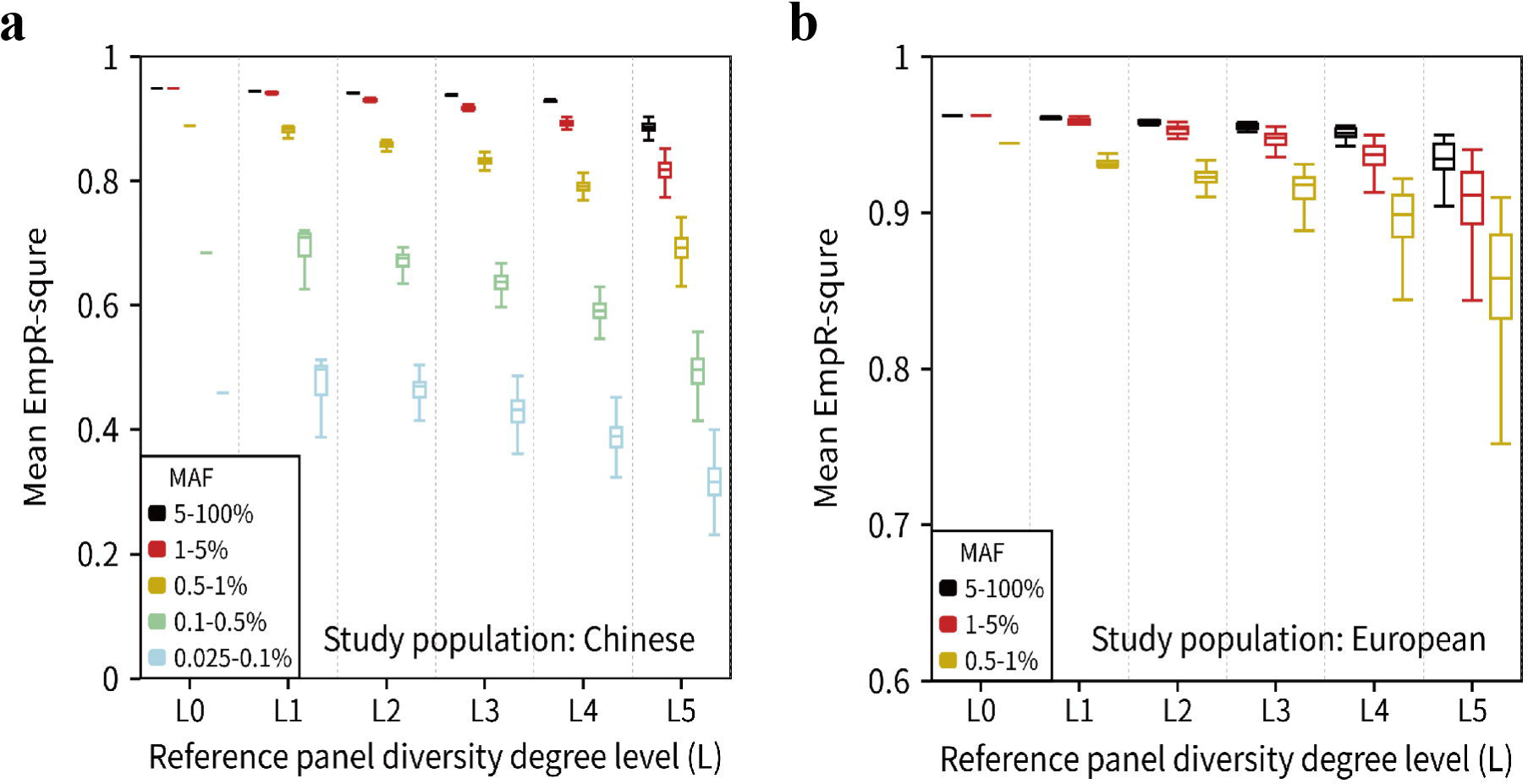
Imputation accuracy for the reference panels with different diversity degrees. **(a)** The boxplot of the EmpR^2^ for Han Chinese using the reference panels with different proportion of EAS populations. Since our study data was Han Chinese population, we used the proportions of non-EAS samples of 1000G (∼0 to 100%) represented the diversity degrees (level_0_ to level_5_) to the GWAS data. All variants were divided into 5 MAF bins. The outliers (mean EmpR^2^ more than Q_3_+1.5*IQR or less than Q_1_-1.5*IQR, IQR=Q_3_-Q_1_) were not plotted. This plot was based on the 126 reference panels, the diversity degree level_0_, level_1_, level_2_, level_3_, level_4_ and level_5_ groups respectively contained 1, 4, 10, 20, 35 and 56 reference panels. **(b)** The boxplot of the EmpR^2^ for European using the reference panels with different proportion of non-EUR populations. Similar to (a), but only three MAF bins of EmpR^2^ of variants were plotted since the variants with MAF < 0.5% were not available (see Methods).

### Imputation accuracy versus size and diversity of panel

We knew that the haplotypes size and populations diversity of reference panel were two crucial factors that affect imputation accuracy. From the results above, we showed that genotype imputation accuracy of rare variants could benefits from the increasing of sample size and an appropriate proportion of diversity of reference panels, respectively. Most of the time, expanding sample size meant to introduce more diverse populations. In this part, we designed a series panel that the haplotypes size and populations diversity simultaneously raised for 85 times. A total of 172 panels were customized and divided into two groups (Figure 1c), sample size expanding from 103 to 868 for Han Chinese group (99 to 864 for European group) while population diversity augmented from 0 to 88%. We found different patterns for Han Chinese and European imputation accuracy. For Han Chinese, the overall accuracy had an improvement at the beginning and reached the highest when panel’s samples increased 3 step-size (27 individuals) with 21% population diversity introduced (Figure 5a). After that, the imputation accuracy continually decreased with diverse populations raised, but still higher than initial panel (0-step panel). However, for the European, the imputation accuracy showed a constantly increased trend from step one to the end with the sample size and diversity grew, the increases from the first step was most obviously (Figure 5a). This result suggested that the positive effect of sample-size-expanding on imputation accuracy was not large enough to neutralize the negative effect of the panel diversity which introduced for Han Chinese after the diversity rare over 21%. But it could offset the negative effect for European and improve the imputation accuracy, which meant that the imputation accuracy for European population could always benefit from the larger panel.

**Figure 5.**
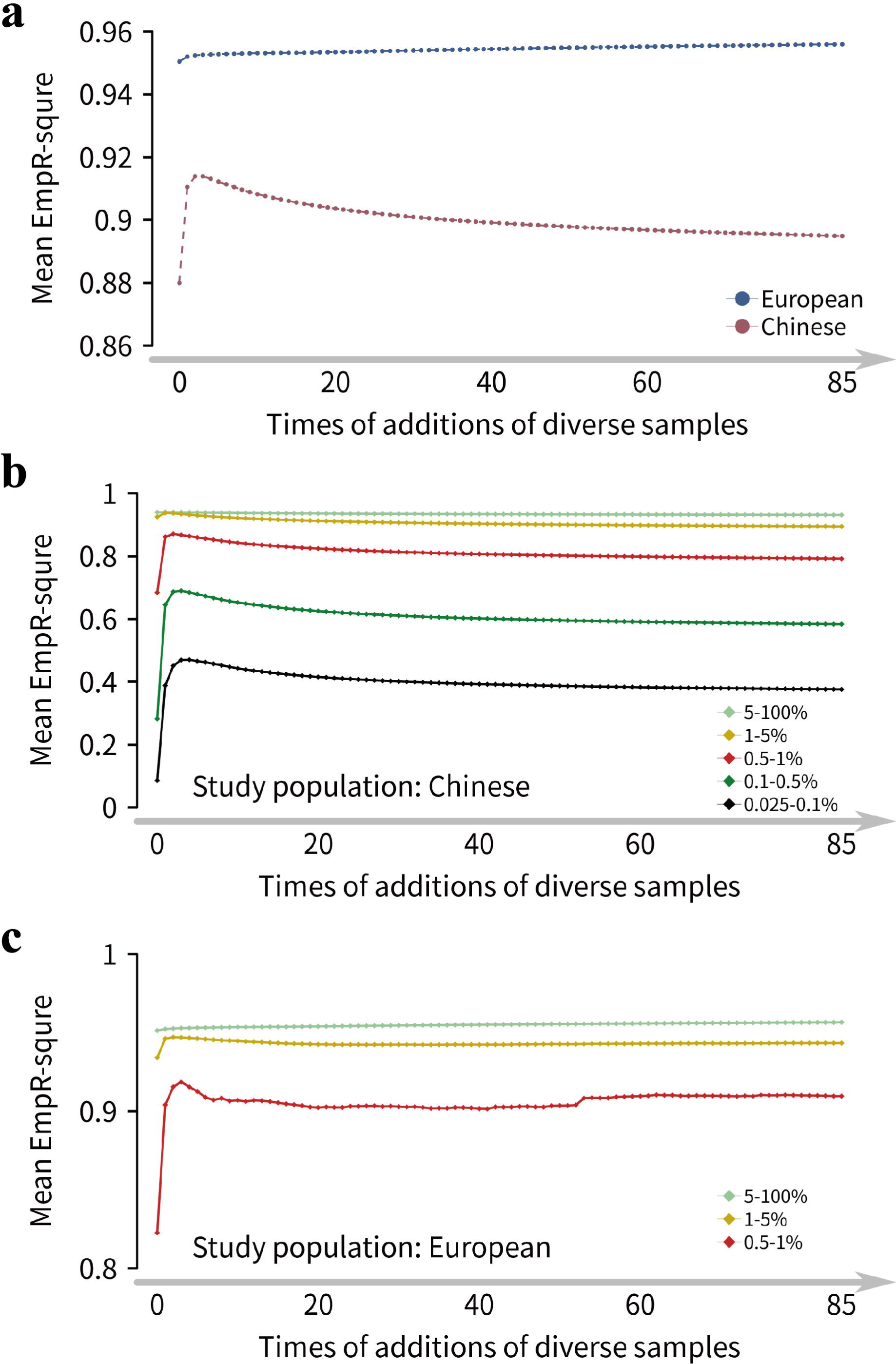
Imputation accuracy for the reference panels with population diversity and sample size constantly increased. **(a)** Overall imputation accuracy trends for Han Chinese and European populations. For Han Chinese, the basic panel (0-step of additions) was CHB (0% diversity, 103 samples with Han Chinese ancestry), the final reference panel was 85-step of additions panel (88% diversity, 868 samples). For the European, the basic panel was CEU (0% diversity, 99 samples with European ancestry). The final reference panel was 85-step of additions panel (88% diversity, 864 samples). **(b)** Imputation accuracy trend for Han Chinese in 5 MAF bins using the reference panels with population diversity and sample size constantly increased. **(c)** Imputation accuracy trend for European, and only three MAF bins of EmpR^2^ of variants were plotted since the variants with MAF < 0.5% were not available (see Methods).

In the divided MAF bins of imputed-variants, the results showed the more detailed changes mode. The imputation accuracy increased rapidly when the diverse population were introduced at the beginning for both populations, and then slowly decreased for the variants of Han Chinese population (Figure 5b) but slowly increased for the common variants of European population (Figure 5c) with diverse samples raised. Besides, for the variants with different MAF bins to reach its highest accuracy, the diversity rate was increased when the variants got more and more rare. These results suggested that an extra diversity of reference panel could remarkably improve the imputation accuracy of rare variants, and appropriate proportion (15% ∼ 21%) diversity could maximize it for Han Chinese population. Besides, we observed that the European population could be more accurately imputed than the Han Chinese.

Based on the design of this part, we developed a panel re-construction tool for researchers to investigate the imputation accuracy in other populations of 1000G. User can set their own study population, diverse populations, step size and adding times in an easily way, and customize a series reference panel. This tool/package could be downloaded now from GitHub, and the details of how to use this tool were included (https://github.com/Abyss-bai/reference-panel-reconstruction).

### Imputation evaluation for the 1000G and HRC and CONVERGE panels

Before actual imputation, the SNPs overlapping between three panels and distribution with seven MAF bins were investigated (Supplementary Figure 3). The Venn diagram showed that 1000G P3, HRC and CONVERGE has 15,524,045, 10,612,366 and 10,550,308 unique sites on autosomes respectively, and all shared 10,303,072 sites in total (Supplementary Figure 4). The HRC-based imputation was performed in Michigan server and the CONVERGE-based imputation was performed in local server, the 1000G-based imputation was conducted in both ways (local and remote), we compared the results of 1000G-based imputation of two servers, the mean EmpR^2^ and imputed sites counts showed perfect consistency (Supplementary Figure 5 and Supplementary Table 3). This result illustrated that the bias caused by local and Michigan imputation server’s difference was negligible. Besides these three reference panels, in this part, we also presented the “CHB21D” panel that consisted of CHB population and 21% extra diverse samples (i.e. the 3-step panel for Han Chinese in last section) to compare its performance with the three big reference panels.

For Han Chinese GWAS sets, the 1000G panel imputed 7,168,371 sites with R^2^ >= 0.8, which was the best. The HRC panel showed the highest imputation quality with mean R^2^ = 0.72 in shared sites and the CONVERGE panel showed the highest imputation accuracy with EmpR^2^ = 0.92. However, due to only about 22 million sites in total were contained in the CONVERGE panel, its number of well-imputed sites was the minimum (5,626,185). For European GWAS sets, the HRC panel resulted in 12,871,067 well-imputed (R^2^ >= 0.8) sites which was the best among the three panels and accounted for 32.9% of total imputed sites. And the HRC panel showed the highest imputation quality with mean R^2^ = 0.73 in 10,302,818 shared imputed sites, and showed the highest imputation accuracy with a quite strong mean EmpR^2^ (0.98) (Table 1). We could clearly know that HRC was the best panel for European samples genotype imputation from these results, but for Chinese samples, however, each of three panels had their own advantages and none of them was the most suitable panel.

We also divided imputed variants by MAF bins as above, but not focused on >5% MAF bin. For Han Chinese GWAS sets, the absolute number of well-imputed SNPs of the HRC and 1000G panels was close and the most, the CONVERGE panel showed the minimum in four MAF bins, and even lower than CHB21D panel (Figure 6a). The average R^2^ which represents imputation quality of variants showed that the CONVERGE panel was slightly better than 1000G panel while the HRC was still the best (Supplementary Figure 6a). For European GWAS sets, the absolute number of well-imputed variants and mean R^2^ of the HRC panel in all four MAF bins were obviously higher than the corresponding values of the 1000G and CONVERGE panels (Figure 6b and Supplementary Figure 6b). Moreover, even for the very rare variants whose MAF in 0.025∼0.1% bin, the HRC panel could well impute about 2.4 million sites while the 1000G and CONVERGE panels could orefnly well impute 0.4 million and 2,021 sites respectively.

**Figure 6.**
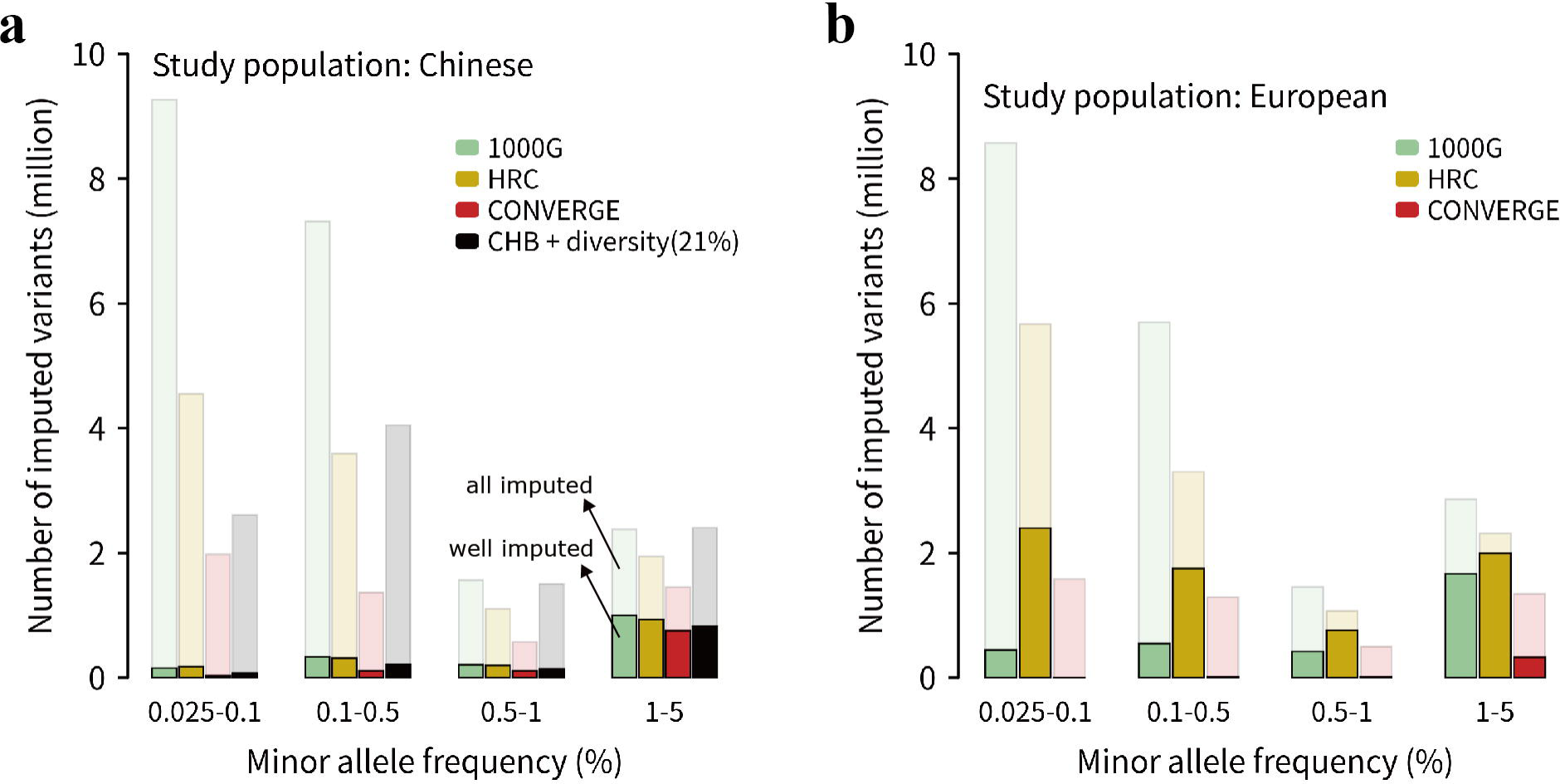
Number of imputed variants for the 1000G, HRC and CONVERGE reference panels. **(a)** Imputation accuracy for the Han Chinese using four different reference panels. Only the mean EmpR^2^ of low-frequency and rare variants were plotted, and the variants were divided into 4 MAF bins. The panel of “CHB + diversity (21%)” was refer to CHB21D panel, which consisted of CHB population and 21% extra diverse samples (i.e. the 3-step panel for Han Chinese in last section). **(b)** Imputation accuracy for the European using three reference panels, and only three MAF bins of EmpR^2^ of variants were plotted since the variants with MAF < 0.5% were not available (see Methods)

To obtain more comparable results, we extracted the 10,302,818 shared sites by three panels. The HRC panel also showed the largest number of well-imputed variants and the highest average of R^2^ in four MAF bins for both populations (Supplementary Figure 7). Besides, we found that, for Han Chinese population, although the HRC imputed the most variants with R^2^ > 0.8, its EmpR^2^ was just the lowest among four panels (Figure 7a). The Chinese-specific panel, CONVERGE, has the highest imputation accuracy, and followed by the CHB21D panel which only contained 130 samples. The constitution of CHB21D panel showed the great potential at imputation accuracy. For European imputation accuracy, the EmpR^2^ of the HRC panel was slightly higher than the 1000G panel, the CONVERGE panel was lowest (Figure 7b). These results implied that, for the European population imputation, the HRC panel was the best choice, for the Chinese population, a high quality and decent reference panel was still needed.

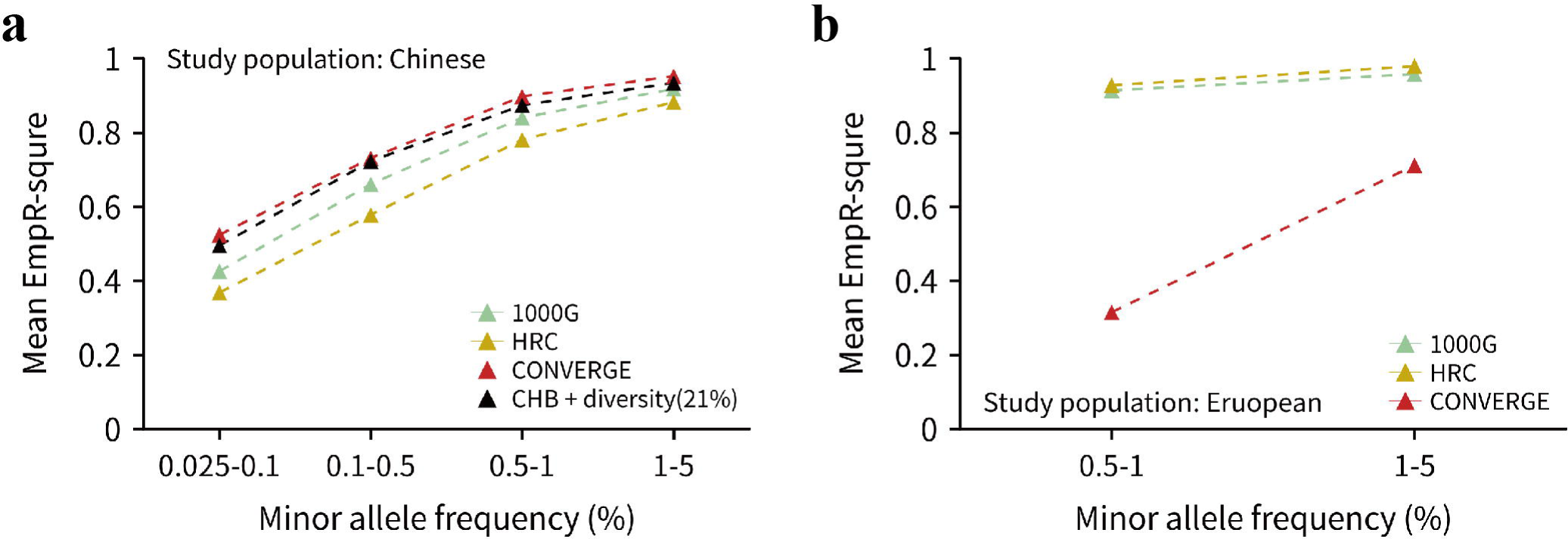

## Discussion

In this study, by conducting 622 imputations with 394 customized reference panels for Han Chinese and European populations, we found that when the haplotypes size of reference panel was increased with extra population-diverse samples, which was usually the real case, the imputation accuracy of Chinese population could reach the highest when reference panel contains a fraction of extra diverse sample (15%∼21%), but when the size of reference panel was fixed, the pattern was different. In addition, we first evaluated the performance of Chinese-specific panel CONVERGE and two frequently used reference panel HRC and 1000G. No doubt the HRC was the best panel for European population since its performance on both imputation accuracy and quality outperformed other panels. For Han Chinese population, the performance of the HRC and 1000G reference panel on well-imputed variants number were better than the CONVERGE panel, but the CONVERGE showed the highest imputation accuracy. However, large-scale Han population reference panel with high quality is remaining needed.

Since the first GWAS published in 2005, scientists showed a fantastic enthusiasm in this powerful design to investigate the genetic risk factors for complex traits. A total of ∼3,700 GWAS in complex trait/diseases have identified thousands of risk variants over the past 14 years[36], and GWAS will continue to contribute knowledge about population genetics in future. The cost of one-human-genome sequencing has dropped to 1000 dollars in 2017[37] and even lower by now, however, it is still too expensive to sequence samples in a large-scale study. With the accomplishment and establishment of large sequencing projects and GWAS databases, such as the 1000G project and UK biobank (UKB), more and more large-scale and high-depth sequencing data went public and available. With reference panels that build from these high quality data, such as HRC, genotype imputation delivered an attractive low-cost alternative to sequencing.

Previous efforts have been focused on imputation evaluation for different populations, such as African, Chinese and Southeast Asian, by using publicly available reference panel[7, 8, 29], despite all these efforts, most studies have been conducted in a relatively shallow way because of the tremendous pressure to the computation server, and the main purpose of these studies were to evaluate the percentage of well imputed SNPs for a suitable reference panel. Less of them investigated how the factors affect the accuracy. Huang and Li’s study designed a series of size-unfixed reference panels using 210 HapMap samples which consist of four populations, and concluded that a mixed panel could lead to the maximal imputation accuracy for a particular population as its primary component was the same HapMap reference panel[19]. Their works investigated how to maximize the accuracy with HapMap reference panel, and raised the importance of size and composition of the reference panel, but was not detailed enough for a systemic study on imputation accuracy of rare variants. The genotype imputation required a high computing power, and the large-scale imputation study was mostly hindered by this requirement.

In this study, we randomly selected 2,000 unrelated individuals for the Han Chinese and the European GWAS set, respectively. Actually, the GWAS set sample size would linearly increase the computed pressure. Usually, for a large-scale imputation accuracy study, people would prefer a smaller GWAS sample size, such as less than 1,000. Although the sample size we used would produce more computation load, it can do reduce the accuracy error and result a more precise and reliable result. The QC we performed was quite strict, besides the common QC steps of imputation (control the high missingness rate of samples and variants, high deviations from Hardy-Weinberg equilibrium and high inbreeding coefficient etc), we checked our GWAS data twice in case of the mistake that varied situations may bring in subsequent imputation, such as SNP mismatch, allele switch and strand flip, by Michigan Imputation Server and local tools. We believed that the GWAS data with a high quality could leads to the results with a high accuracy. Note that our European GWAS data had been QCed before, the variants with MAF < 0.5% had been removed, which meant that the imputation performance on rare variants of European population were not available. However, we included all variants in Han Chinese GWAS data, even for singletons. The common variants can be accurately imputed by any existing big reference panel, such as the 1000G and the HRC panel. We used the rare variants set in Han Chinese to study the genotyping imputation accuracy changes patterns with different composition of reference panels.

All the customized reference panels in this study were based on the 1000G Phase 3, there were two popular versions for the 1000G panel, one included singletons and another did not. We found that the Sanger Imputation Server used singleton-included version and the Michigan Imputation Server used non-singleton version. We investigated the imputation performance of these two versions of panel on Han Chinese data in local and found that the difference between them were negligible (Supplementary Figure 5 and Supplementary Table 3). After all, we decided to use the non-singleton version and were consistent with Michigan server since the imputation tool that we employed were both Minimac3. The customized reference panels fell into three categories: 1) haplotype size changed and the population diversity ratio was fixed. 2) Population diversity degree changed and the haplotype size was fixed. 3) Both of them changed with a fixed step-size, note that the increased diversity ratio became more and more small because the panel size got more and more large. In the design of our second category panels, we made a simple cluster analysis by the longitude and latitude of populations, the result showed that almost all populations obviously followed by the 1000G groups (EAS, EUR, AFR, AMR and SAS), except the ‘ACB’ population (Supplementary Figure 8). The ‘ACB’ was classified into the AFR group in the 1000G, but it was far more close to the AMR group geographically, so we reclassified this population into the AMR group. But in reality, it will not make a nontrivial difference in our study, because our GWAS set were Han Chinese and European population, the ‘ACB’ was always diverse to our study sets. Beside the haplotypes size and diversity of reference panel, the sequencing coverage would also affect the imputation accuracy. The high-depth sequencing could result the more accurate genotypes in the reference panel, which in turn improve the accuracy of the inferred haplotypes[1]. We did not systematically evaluate the influence of sequencing coverage since the rest of effective variables could not be completely controlled by the update public reference panels.

There was a phenomenon that crossed all the imputation results, which is that the imputation performance for the European population was always better than for Han Chinese by their best-panels. Taking the first category panels (described in the previous paragraph, imputation accuracy vs. haplotype size) as an example, the results showed that the best-panel (the CHB panel) for the Han Chinese population got a 0.874 average accuracy while the European got a 0.947 average accuracy by its best-panel (the CEU panel). The first reason that came to mind might cause this gap was that the European GWAS data not included variants with MAF < 0.5%, but the Han Chinese did, even if the variants between them were shared. We then removed all variants with MAF less than 0.5% that in Han Chinese data for both sets, and found the average imputation accuracy raised to 0.929 and 0.948 for the Han Chinese and European respectively, the gap narrowed but still existed. We could know it from the statistic of variants of different MAF bins as well (Figure 2c and 2d). Another reason might cause the gap was the microarray chip, the two GWAS sets used the Illumina 610k BeadChips which was designed for the European population at the beginning.

Another discrepancy of imputation results between the Han Chinese and European population was that, despite the introduced diversity, the accuracy of imputed common variants of European population could always benefit from the expanding haplotypes size of reference panel, while the corresponding accuracy of the Han Chinese population could benefit only when the diversity of reference panel remained a small ratio (Figure 5). This result suggested that, in a sense, the genome of the European may have a higher acceptability than Chinese genome which meant it was more diverse. In the course of evolution, the view of intermarriage of Chinese was more conservative than European[38, 39]. And an open intermarriage view may result in the genome became more and more diverse across generations.

In the last decade, many cohort studies and WGS projects have been conducted, and several genome reference panels were produced. However, most of the these cohorts and reference panels were focused on the European and African American populations, such as the Wellcome Trust Case Control Consortium (WTCCC)[40], UK10K[24], HRC[26] and TOPMed program. Few WGS projects were conducted in Chinese population. The HapMap3 only included 137 native Chinese individuals[20], the 1000G project phase 3 included 301 Chinese individuals, and only 208 were Han Chinese[23]. In 2017, 90 unrelated individuals of Chinese ancestry were sequenced at a high depth (∼80X)[41] by the Beijing Genomics Institute (BGI-Shenzhen). However, the sample size of these WGS projects was small. Although the CONVERGE project sequenced 11,670 female Han Chinese and provided the largest whole genome sequencing resource of Chinese[27], it was only able to call ∼22 million high quality variants and the sequencing coverage of CONVERGE was low (1.7X). At present, we are engaging in a Chinese cohort and have collected 10K samples across 29 provincial regions of China, the sequencing of ∼4000 samples at ∼17X average coverage is ongoing. We hope to generate a high quality and decent population-specific reference panel for public use for the largest ethnic group in the world.

In summary, we systematically investigated the relationship between genotype imputation accuracy of rare variants and the composition of reference panel, and proposed an optimum constituent ratio for the reference panel for the Han Chinese imputation. We found the different patterns of imputation accuracy variation between the European and Han Chinese. This point enlightened us that we should use more special panels when impute the Chinese genome, and this could be generalized to the other populations with conservative genome.

## Acknowledgments

This study was supported by the Zhejiang Provincial Natural Science Foundation for Distinguished Young Scholars of China (LR17H070001) and by the National Natural Science Foundation of China (81871831). The funding agencies had no role in the study design, data collection and analysis, decision to publish or preparation of the manuscript. We thank the peer reviewers for their thorough and helpful review of this manuscript.

---

*the Han Chinese and European populations*

| N | Well-imputed variants number<br>(proportion) | Shared variants |  | *Genotyped variants |  |
| --- | --- | --- | --- | --- | --- |
|  |  | N | Mean R<sup>2</sup> (SD) | N | Mean EmpR<sup>2</sup> (SD) |
| 39,127,690 | 6,228,449 (15.9%) | 10,302,818 | 0.72 (0.19) | 515,754 | 0.89 (0.17) |
| 47,109,465 | 7,168,371 (15.2%) | 10,302,818 | 0.68 (0.22) | 516,408 | 0.91 (0.14) |
| 47,109,431 | 6,275,000 (13.3%) | - | - | 516,408 | 0.91 (0.16) |
| 24,114,249 | 5,626,185 (23.3%) | 10,302,818 | 0.58 (0.20) | 511,715 | 0.92 (0.11) |
| 39,127,690 | 12,871,067 (32.9%) | 10,302,818 | 0.73 (0.12) | 515,754 | 0.98 (0.04) |
| 47,109,465 | 9,422,724 (20.0%) | 10,302,818 | 0.69 (0.18) | 516,408 | 0.96 (0.07) |
| 24,114,249 | 4,539,069 (18.8%) | 10,302,818 | 0.51 (0.18) | 511,715 | 0.82 (0.21) |

## References

1. Das S, Abecasis GR, Browning BL: Genotype Imputation from Large Reference Panels. Annu Rev Genomics Hum Genet 2018, 19:73–96.

2. Mahajan A, Taliun D, Thurner M et al: Fine-mapping type 2 diabetes loci to single-variant resolution using high-density imputation and islet-specific epigenome maps. Nat Genet 2018, 50(11):1505–1513.

3. Zheng HF, Forgetta V, Hsu YH et al: Whole-genome sequencing identifies EN1 as a determinant of bone density and fracture. Nature 2015, 526(7571):112–117.

4. Estrada K, Styrkarsdottir U, Evangelou E et al: Genome-wide meta-analysis identifies 56 bone mineral density loci and reveals 14 loci associated with risk of fracture. Nat Genet 2012, 44(5):491–501.

5. Willer CJ, Sanna S, Jackson AU et al: Newly identified loci that influence lipid concentrations and risk of coronary artery disease. Nat Genet 2008, 40(2):161–169.

6. Anderson CA, Pettersson FH, Barrett JC et al: Evaluating the effects of imputation on the power, coverage, and cost efficiency of genome-wide SNP platforms. Am J Hum Genet 2008, 83(1):112–119.

7. Lin Y, Liu L, Yang S et al: Genotype imputation for Han Chinese population using Haplotype Reference Consortium as reference. Hum Genet 2018, 137(6-7):431–436.

8. Vergara C, Parker MM, Franco L et al: Genotype imputation performance of three reference panels using African ancestry individuals. Hum Genet 2018, 137(4):281–292.

9. Zheng HF, Ladouceur M, Greenwood CM et al: Effect of genome-wide genotyping and reference panels on rare variants imputation. J Genet Genomics 2012, 39(10):545–550.

10. Gibson G: Rare and common variants: twenty arguments. Nat Rev Genet 2012, 13(2):135–145.

11. Barbujani G, Colonna V: Human genome diversity: frequently asked questions. Trends Genet 2010, 26(7):285–295.

12. International Multiple Sclerosis Genetics Consortium. Electronic address ccye, International Multiple Sclerosis Genetics C: Low-Frequency and Rare-Coding Variation Contributes to Multiple Sclerosis Risk. Cell 2018, 175(6):1679–1687 e1677.

13. Tin A, Li Y, Brody JA et al: Large-scale whole-exome sequencing association studies identify rare functional variants influencing serum urate levels. Nat Commun 2018, 9(1):4228.

14. Gazal S, Loh PR, Finucane HK et al: Functional architecture of low-frequency variants studies by imputation of genotypes. Nat Genet 2007, 39(7):906–913.

19. Huang L, Li Y, Singleton AB et al: Genotype-imputation accuracy across worldwide human populations. Am J Hum Genet 2009, 84(2):235–250.

20. International HapMap C, Altshuler DM, Gibbs RA et al: Integrating common and rare genetic variation in diverse human populations. Nature 2010, 467(7311):52–58.

21. Huang J, Howie B, McCarthy S et al: Improved imputation of low-frequency and rare variants using the UK10K haplotype reference panel. Nat Commun 2015, 6:8111.

22. Deelen P, Menelaou A, van Leeuwen EM et al: Improved imputation quality of low-frequency and rare variants in European samples using the ‘Genome of The Netherlands’. Eur J Hum Genet 2014, 22(11):1321–1326.

23. Genomes Project C, Auton A, Brooks LD et al: A global reference for human genetic variation. Nature 2015, 526(7571):68–74.

24. Consortium UK, Walter K, Min JL et al: The UK10K project identifies rare variants in health and disease. Nature 2015, 526(7571):82–90.

25. Boomsma DI, Wijmenga C, Slagboom EP et al: The Genome of the Netherlands: design, and project goals. Eur J Hum Genet 2014, 22(2):221–227.

26. McCarthy S, Das S, Kretzschmar W et al: A reference panel of 64,976 haplotypes for genotype imputation. Nat Genet 2016, 48(10):1279–1283.

27. Cai N, Bigdeli TB, Kretzschmar WW et al: 11,670 whole-genome sequences representative of the Han Chinese population from the CONVERGE project. Sci Data 2017, 4:170011.

28. Nelson SC, Stilp AM, Papanicolaou GJ et al: Improved imputation accuracy in Hispanic/Latino populations with larger and more diverse reference panels: applications in the Hispanic Community Health Study/Study of Latinos (HCHS/SOL). Hum Mol Genet 2016, 25(15):3245–3254.

29. Lert-Itthiporn W, Suktitipat B, Grove H et al: Validation of genotype imputation in Southeast Asian populations and the effect of single nucleotide polymorphism annotation on imputation outcome. BMC Med Genet 2018, 19(1):23.

30. Jostins L, Morley KI, Barrett JC: Imputation of low-frequency variants using the HapMap3 benefits from large, diverse reference sets. Eur J Hum Genet 2011, 19(6):662–666.

31. Han JW, Zheng HF, Cui Y et al: Genome-wide association study in a Chinese Han population identifies nine new susceptibility loci for systemic lupus erythematosus. Nat Genet 2009, 41(11):1234–1237.

37. Shendure J, Balasubramanian S, Church GM et al: DNA sequencing at 40: past, present and future. Nature 2017, 550(7676):345–353.

38. Jian Z: The recent trend of ethnic intermarriage in China: an analysis based on the census data. The Journal of Chinese Sociology 2017, 4(1):11.

39. Alba RD: Intermarriage and ethnicity among European Americans. Contemporary Jewry 1991, 12(1):3–19.

40. Wellcome Trust Case Control C, Craddock N, Hurles ME et al: Genome-wide association study of CNVs in 16,000 cases of eight common diseases and 3,000 shared controls. Nature 2010, 464(7289):713–720.

41. Lan T, Lin H, Zhu W et al: Deep whole-genome sequencing of 90 Han Chinese genomes. Gigascience 2017, 6(9):1–7.

