## Supplemental Figures for "Genotype Imputation and Reference Panel: A Systematic Evaluation"

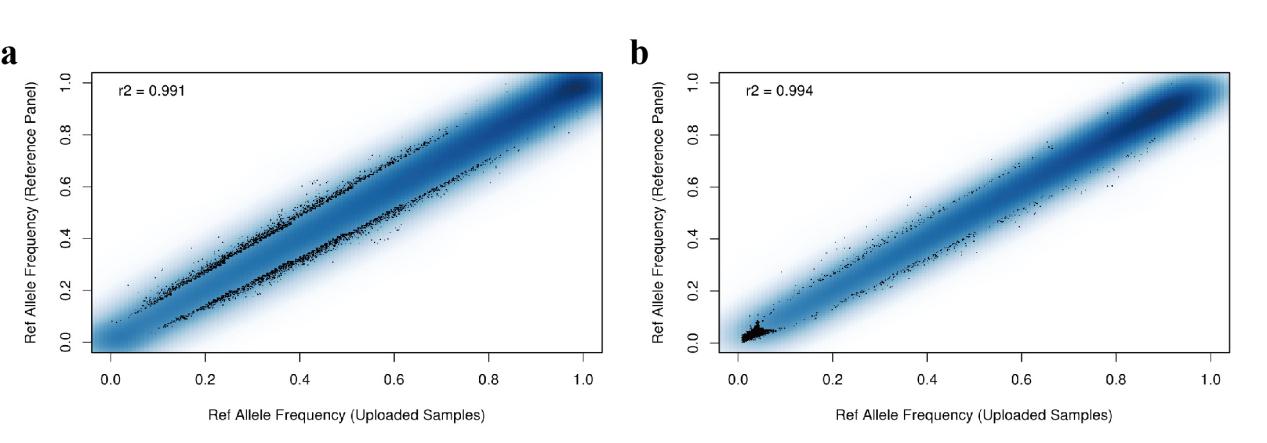

**Supplementary Figure 1. Allele frequency correlation plot between study samples and 1000G reference panel**

This plot was from the allele frequency QC report of study data by Michigan Imputation Server. The plot showed the densities of frequencies falling into each part (excluding chromosome X). The first 5000 points from areas of lowest regional densities were plotted. The reference panels were **(a)** EAS and **(b)** EUR of the 1000G Phase3, corresponding to the Han Chinese and European populations respectively. We can see that the allele frequency of variants distribution of Han Chinese data was more uniform than the European data. And the density of low-frequency variants of the European data was lower than Han Chinese data.

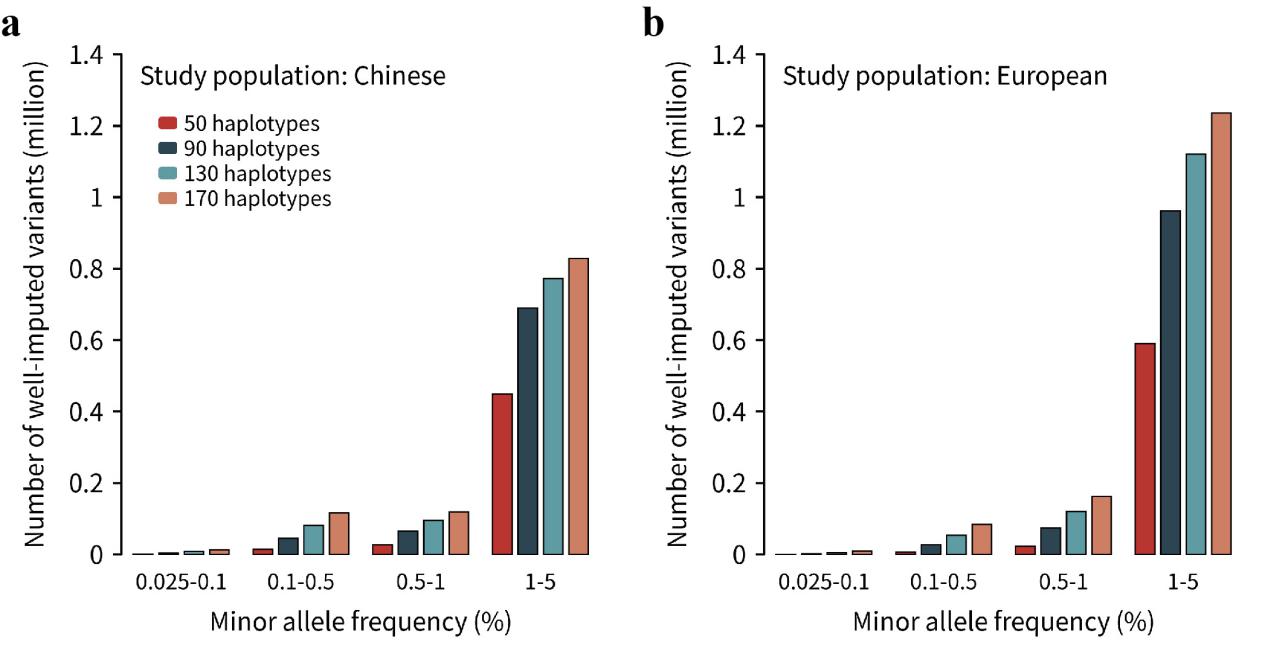

**Supplementary Figure 2. Number of well-imputed variants for the reference panels with four haplotype size gradients.**

The number of imputed variants with mean R2 >0.8 in four MAF bins for **(a)** the Han Chinese and **(b)** European populations data.

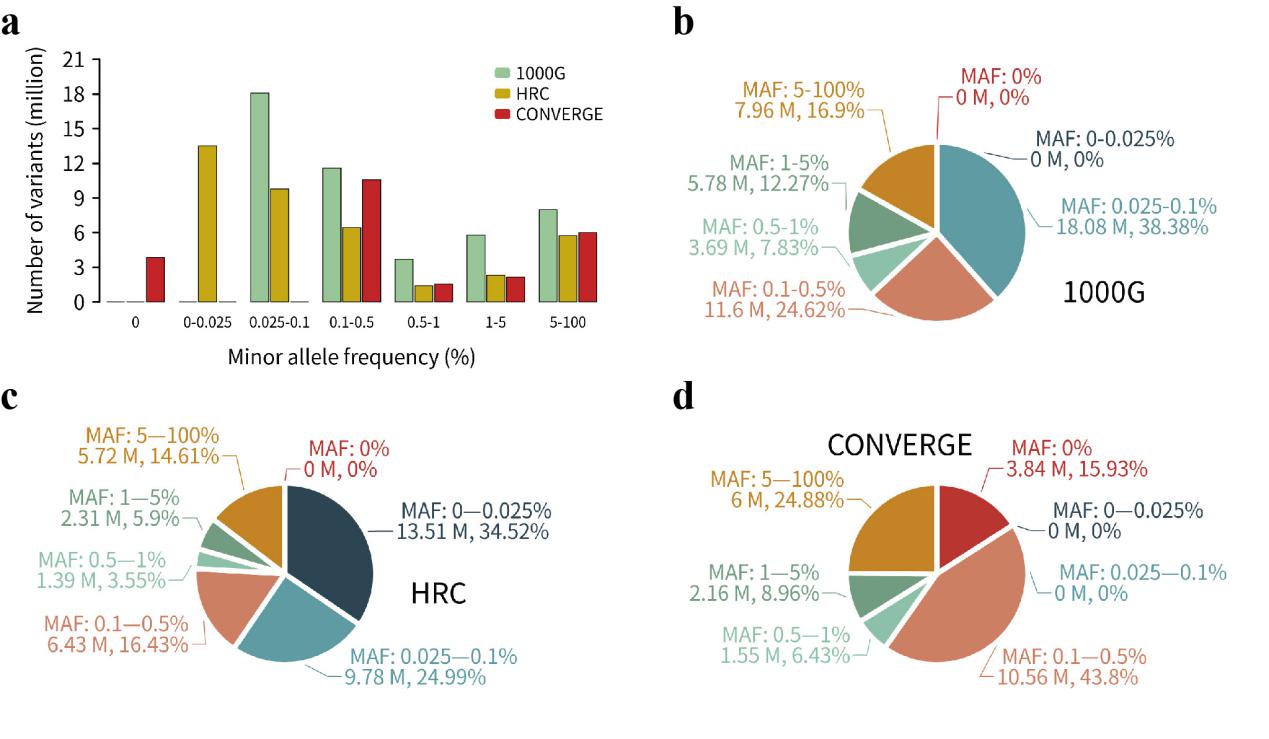

**Supplementary Figure 3. Summary statistics of the 1000G, HRC and CONVERGE reference panels**

The 1000G reference panel we used was the no singleton version. **(a)** All variants were divided into 7 MAF bins for the detailed comparison between the three panels. **(b)** The 1000G Phase3 reference panel. **(c)** The HRC reference panel. **(d)** The CONVERGE reference panel.

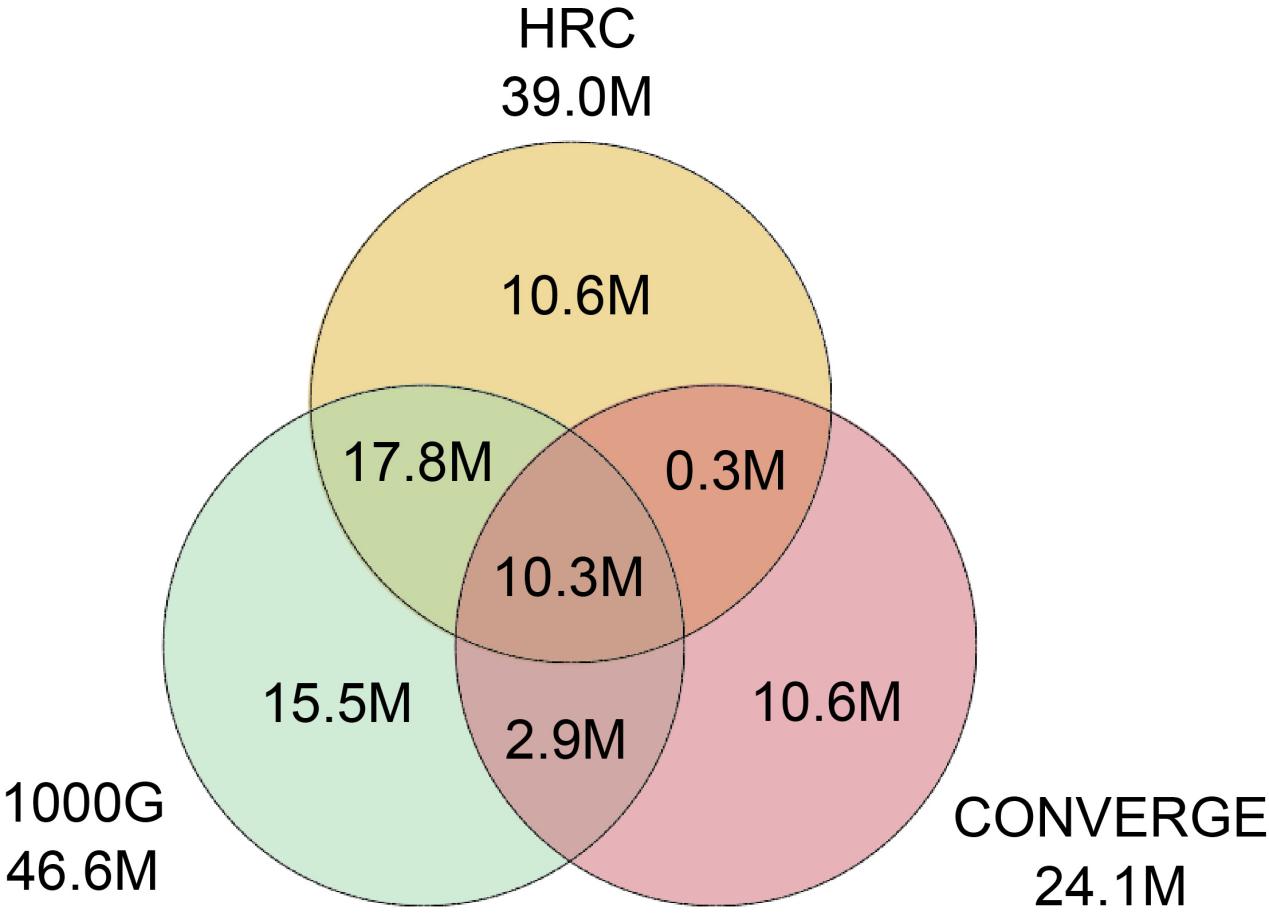

**Supplementary Figure 4. The Venn diagram of imputed variants of autosomes for the 1000G, HRC and CONVERGE reference panels.**

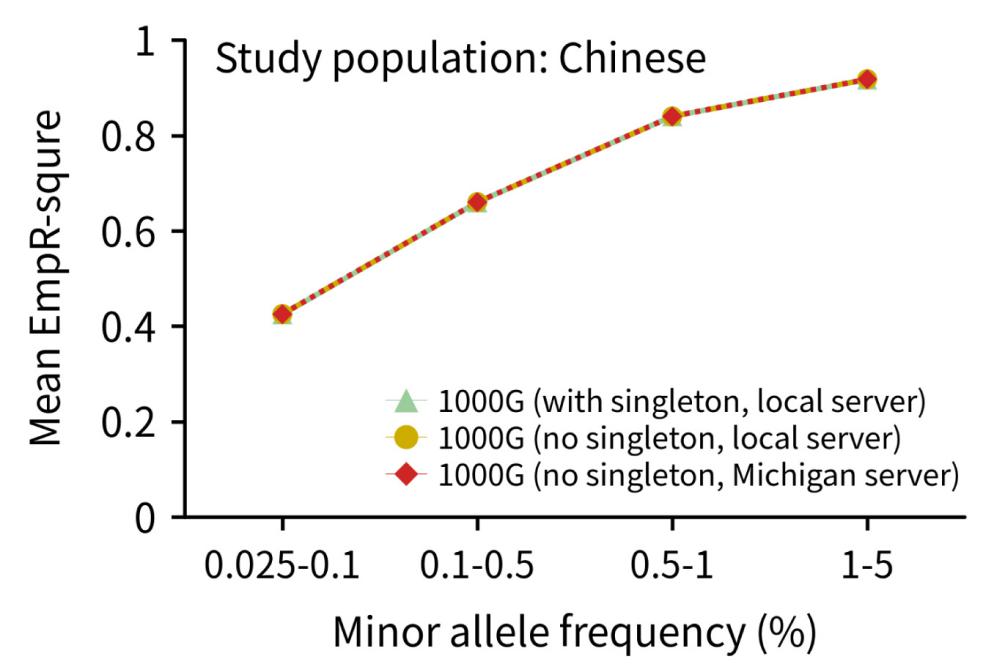

**Supplementary Figure 5. Imputation accuracy for different imputation strategies.**

These three imputations were performed for the results consistency test between local and remote (Michigan) imputation server using 1000G reference panels with two versions (with singleton or not). The imputation accuracy for the Han Chinese population in 4 MAF bins was plotted.

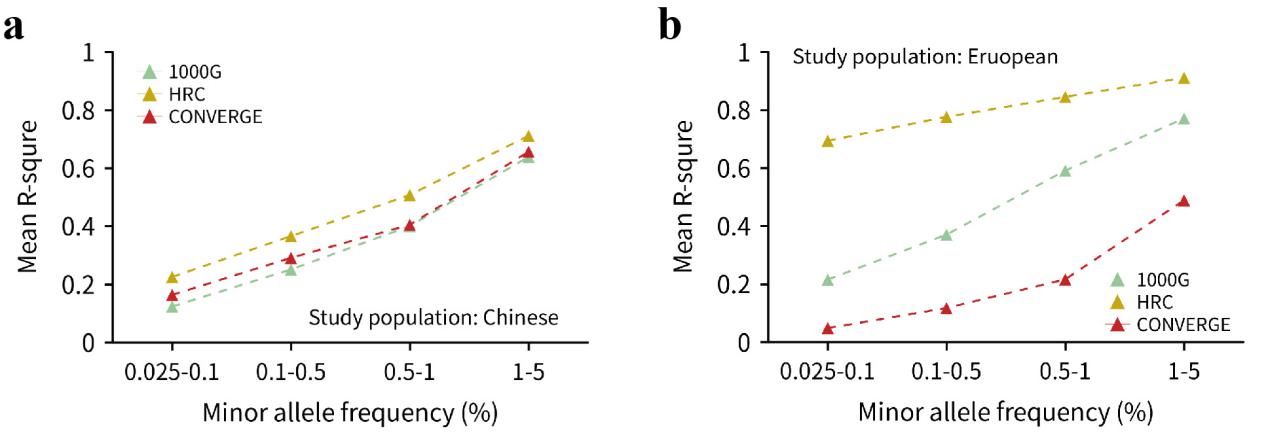

**Supplementary Figure 6. Imputation quality for the 1000G, HRC and CONVERGE reference panels**

The mean R^2^ (measuring the imputation quality) of all imputed variants of four reference panels for **(a)** Han Chinese and three panels for **(b)** European populations. The common variants were not focused because of its reliability of imputation. All variants with MAF <5% were divided into 4 MAF bins.

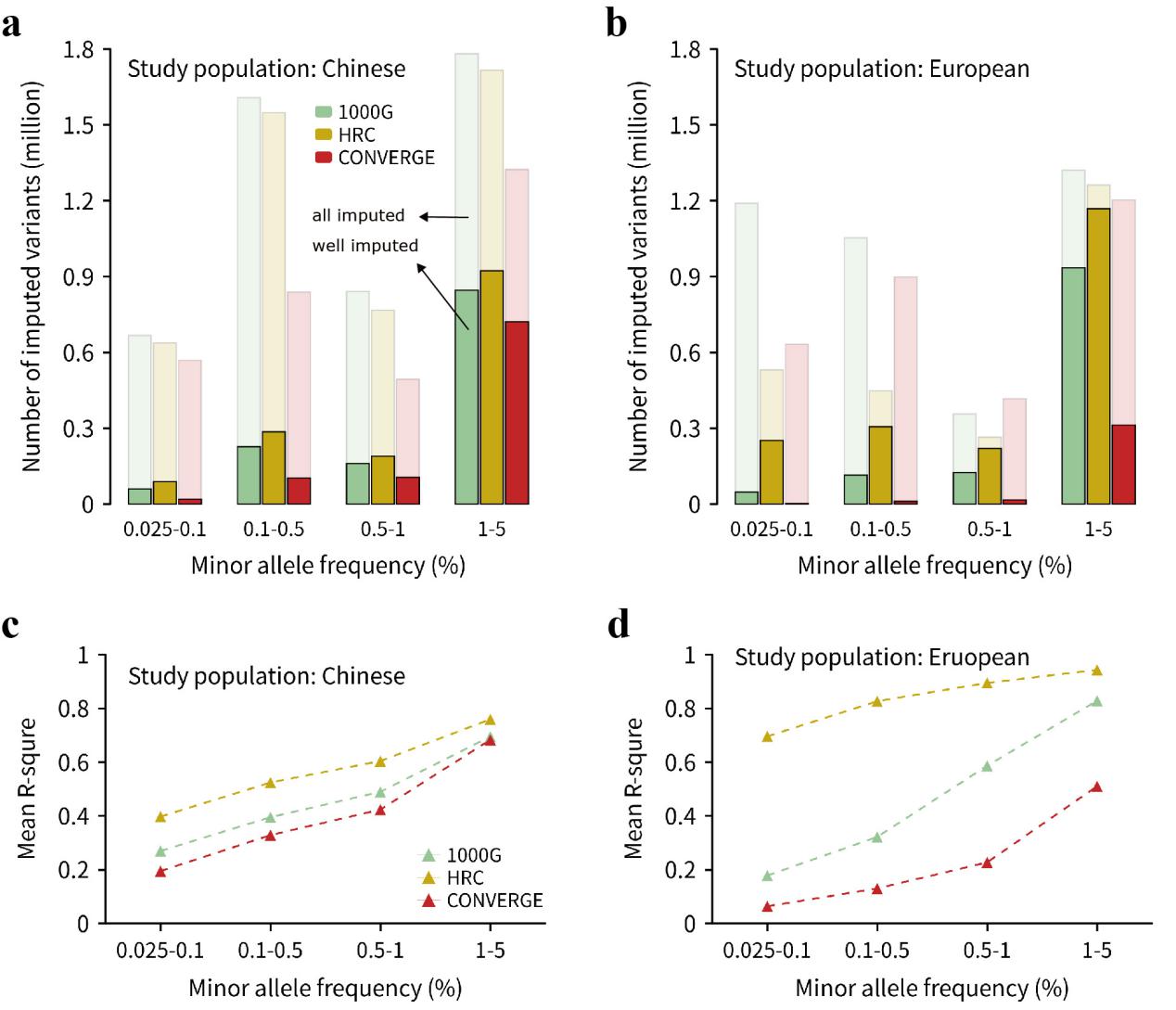

**Supplementary Figure 7. Imputation performance of shared variants for the 1000G, HRC and CONVERGE reference panels**

The number of shared imputed variants of the three reference panels for **(a)** Han Chinese and **(b)** European populations. All variants with MAF <5% were divided into 4 MAF bins. The different colors represented different reference panels, and the light color represented all imputed variants, the dark color represented well imputed variants (R^2^ >0.8). The mean R^2^ of 4 different MAF bins was plotted for **(c)** the Han Chinese and **(d)** European populations.

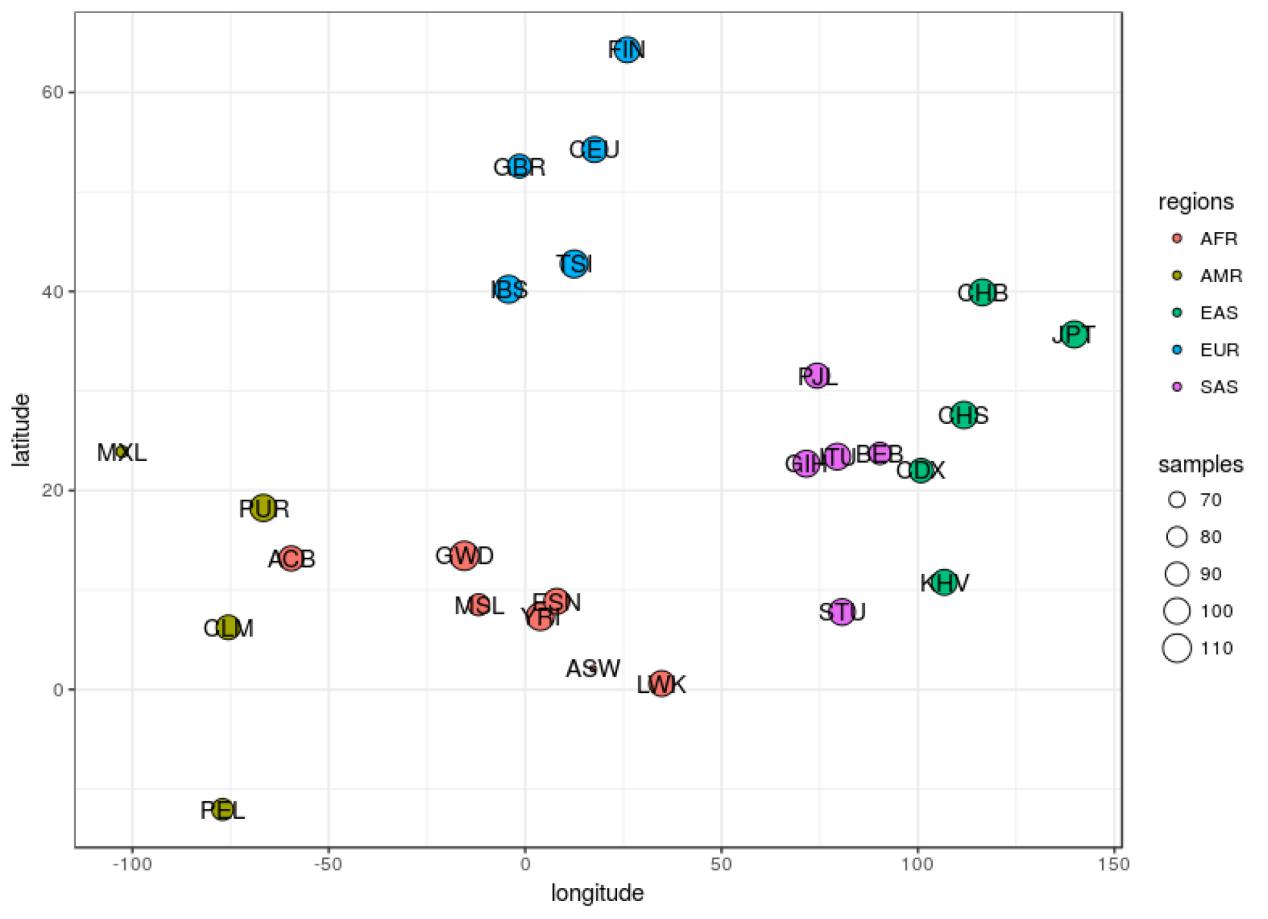

**Supplementary Figure 8. Geographically cluster plot for 25 populations of the 1000G Phase3**

The ACB population belongs to AFR group in the 1000G, and it is close to AMR group in geographically. However, it was always diverse to our study populations (the Han Chinese and European).

***Supplementary Table 1: The composition of 5 Groups extracted from the 1000G Phase 3***

| Groups 1 to 5 | Populations 1 to 5 | Haplotype Number |
| --- | --- | --- |
| EAS | CHB | 128 |
|  | CHS | 128 |
|  | CDX | 128 |
|  | JPT | 128 |
|  | KHV | 128 |
| EUR | CEU | 128 |
|  | TSI | 128 |
|  | FIN | 128 |
|  | GBR | 128 |
|  | IBS | 128 |
| AFR | YRI | 128 |
|  | LWK | 128 |
|  | GWD | 128 |
|  | MSL | 128 |
|  | ESN | 128 |
| AMR | PUR | 128 |
|  | *ACB | 128 |
|  | CLM | 128 |
|  | PEL | 128 |
|  | MXL | 128 |
| SAS | GIH | 128 |
|  | PJL | 128 |
|  | BEB | 128 |
|  | STU | 128 |
|  | ITU | 128 |

* This population belongs to AFR group in the 1000G Phase 3. All samples were randomly selected.

***Supplementary Table 2: The composition of all 126 diverse reference panels***

Please check the Supplement-Table-2.xlsx file.

***Supplementary Table 3: 1000G reference panels imputation performance in Han Chinese with different conditions***

| panel | MafBin | SNPNum | EmpR^2^>=0.8 | EmpR^2^<0.8 | Mean EmpR^2^ | Median EmpR^2^ | EmpR^2^ SD |
| --- | --- | --- | --- | --- | --- | --- | --- |
| 1000G panel without singleton and performed in Michigan server | 0 | 8 | 0 | 8 | 0 | 0 | 0 |
|  | 0-0.025% | 942 | 189 | 753 | 0.24964 | 2.00E-05 | 0.39211 |
|  | 0.025-0.1% | 12361 | 2510 | 9851 | 0.42574 | 0.34976 | 0.33129 |
|  | 0.1-0.5% | 26210 | 10919 | 15291 | 0.6606 | 0.72636 | 0.29101 |
|  | 0.5-1% | 8008 | 5871 | 2137 | 0.84033 | 0.93399 | 0.21504 |
|  | 1-5% | 42132 | 36956 | 5176 | 0.91814 | 0.97604 | 0.14175 |
|  | 5-100% | 426747 | 393203 | 33544 | 0.94064 | 0.98194 | 0.10939 |
| 1000G panel without singleton and performed in local server | 0 | 8 | 0 | 8 | 0 | 0 | 0 |
|  | 0-0.025% | 942 | 189 | 753 | 0.24965 | 2.00E-05 | 0.39212 |
|  | 0.025-0.1% | 12362 | 2513 | 9849 | 0.42577 | 0.34983 | 0.33129 |
|  | 0.1-0.5% | 26209 | 10920 | 15289 | 0.6606 | 0.72631 | 0.29101 |
|  | 0.5-1% | 8008 | 5871 | 2137 | 0.84033 | 0.93399 | 0.21505 |
|  | 1-5% | 42132 | 36958 | 5174 | 0.91814 | 0.97602 | 0.14175 |
|  | 5-100% | 426747 | 393214 | 33533 | 0.94064 | 0.98194 | 0.10939 |
| 1000G panel with singleton and performed in local server | 0 | 9 | 0 | 9 | 0 | 0 | 0 |
|  | 0-0.025% | 932 | 186 | 746 | 0.24789 | 2.00E-05 | 0.39216 |
|  | 0.025-0.1% | 12373 | 2514 | 9859 | 0.42509 | 0.34896 | 0.3313 |
|  | 0.1-0.5% | 26212 | 10922 | 15290 | 0.66008 | 0.72474 | 0.2913 |
|  | 0.5-1% | 8001 | 5868 | 2133 | 0.84064 | 0.93371 | 0.21448 |
|  | 1-5% | 42143 | 36928 | 5215 | 0.91766 | 0.97595 | 0.14278 |
|  | 5-100% | 426740 | 392962 | 33778 | 0.94032 | 0.9819 | 0.11007 |

EmpR^2^ means empirical-R^2^, which was used to measure the imputation accuracy using the variants that were genotyped by the Illumina Human610-Quad (610K) BeadChip.. All variants were divided into 7 MAF bins for the detailed comparison between three strategies.
