## Supplemental Tables for "Genotype Imputation and Reference Panel: A Systematic Evaluation"

***Supplementary Table 1: The composition of 5 Groups extracted from the 1000G Phase 3***

| Groups 1 to 5 | Populations 1 to 5 | Haplotype Number |
| --- | --- | --- |
| EAS | CHB | 128 |
|  | CHS | 128 |
|  | CDX | 128 |
|  | JPT | 128 |
|  | KHV | 128 |
| EUR | CEU | 128 |
|  | TSI | 128 |
|  | FIN | 128 |
|  | GBR | 128 |
|  | IBS | 128 |
| AFR | YRI | 128 |
|  | LWK | 128 |
|  | GWD | 128 |
|  | MSL | 128 |
|  | ESN | 128 |
| AMR | PUR | 128 |
|  | *ACB | 128 |
|  | CLM | 128 |
|  | PEL | 128 |
|  | MXL | 128 |
| SAS | GIH | 128 |
|  | PJL | 128 |
|  | BEB | 128 |
|  | STU | 128 |
|  | ITU | 128 |

* This population belongs to AFR group in the 1000G Phase 3. All samples were randomly selected.

***Supplementary Table 2: The composition of all 126 diverse reference panels***

| Index  of RP | Combinations  of 5 Groups | Pop. of EAS | Pop. of EUR | Pop. of AFR | Pop. of AMR | Pop. of SAS |
| --- | --- | --- | --- | --- | --- | --- |
| 1 | (5, 0, 0, 0, 0) | CHB,CHS,CDX,JPT,KHV | - | - | - | - |
| 2 | (4, 1, 0, 0, 0) | CHB,CHS,CDX,JPT | CEU | - | - | - |
| 3 | (4, 0, 1, 0, 0) | CHB,CHS,CDX,JPT | - | YRI | - | - |
| 4 | (4, 0, 0, 1, 0) | CHB,CHS,CDX,JPT | - | - | PUR | - |
| 5 | (4, 0, 0, 0, 1) | CHB,CHS,CDX,JPT | - | - | - | GIH |
| 6 | (3, 2, 0, 0, 0) | CHB,CHS,CDX | CEU,TSI | - | - | - |
| 7 | (3, 1, 1, 0, 0) | CHB,CHS,CDX | CEU | YRI | - | - |
| 8 | (3, 1, 0, 1, 0) | CHB,CHS,CDX | CEU | - | PUR | - |
| 9 | (3, 1, 0, 0, 1) | CHB,CHS,CDX | CEU | - | - | GIH |
| 10 | (3, 0, 2, 0, 0) | CHB,CHS,CDX | - | YRI,LWK | - | - |
| 11 | (3, 0, 1, 1, 0) | CHB,CHS,CDX | - | YRI | PUR | - |
| 12 | (3, 0, 1, 0, 1) | CHB,CHS,CDX | - | YRI | - | GIH |
| 13 | (3, 0, 0, 2, 0) | CHB,CHS,CDX | - | - | PUR,ACB | - |
| 14 | (3, 0, 0, 1, 1) | CHB,CHS,CDX | - | - | PUR | GIH |
| 15 | (3, 0, 0, 0, 2) | CHB,CHS,CDX | - | - | - | GIH,PJL |
| 16 | (2, 3, 0, 0, 0) | CHB,CHS | CEU,TSI,FIN | - | - | - |
| 17 | (2, 2, 1, 0, 0) | CHB,CHS | CEU,TSI | YRI | - | - |
| 18 | (2, 2, 0, 1, 0) | CHB,CHS | CEU,TSI | - | PUR | - |
| 19 | (2, 2, 0, 0, 1) | CHB,CHS | CEU,TSI | - | - | GIH |
| 20 | (2, 1, 2, 0, 0) | CHB,CHS | CEU | YRI,LWK | - | - |
| 21 | (2, 1, 1, 1, 0) | CHB,CHS | CEU | YRI | PUR | - |
| 22 | (2, 1, 1, 0, 1) | CHB,CHS | CEU | YRI | - | GIH |
| 23 | (2, 1, 0, 2, 0) | CHB,CHS | CEU | - | PUR,ACB | - |
| 24 | (2, 1, 0, 1, 1) | CHB,CHS | CEU | - | PUR | GIH |
| 25 | (2, 1, 0, 0, 2) | CHB,CHS | CEU | - | - | GIH,PJL |
| 26 | (2, 0, 3, 0, 0) | CHB,CHS | - | YRI,LWK,GWD | - | - |
| 27 | (2, 0, 2, 1, 0) | CHB,CHS | - | YRI,LWK | PUR | - |
| 28 | (2, 0, 2, 0, 1) | CHB,CHS | - | YRI,LWK | - | GIH |
| 29 | (2, 0, 1, 2, 0) | CHB,CHS | - | YRI | PUR,ACB | - |
| 30 | (2, 0, 1, 1, 1) | CHB,CHS | - | YRI | PUR | GIH |
| 31 | (2, 0, 1, 0, 2) | CHB,CHS | - | YRI | - | GIH,PJL |
| 32 | (2, 0, 0, 3, 0) | CHB,CHS | - | - | PUR,ACB,CLM | - |
| 33 | (2, 0, 0, 2, 1) | CHB,CHS | - | - | PUR,ACB | GIH |
| 34 | (2, 0, 0, 1, 2) | CHB,CHS | - | - | PUR | GIH,PJL |
| 35 | (2, 0, 0, 0, 3) | CHB,CHS | - | - | - | GIH,PJL,BEB |
| 36 | (1, 4, 0, 0, 0) | CHB | CEU,TSI,FIN,GBR | - | - | - |
| 37 | (1, 3, 1, 0, 0) | CHB | CEU,TSI,FIN | YRI | - | - |
| 38 | (1, 3, 0, 1, 0) | CHB | CEU,TSI,FIN | - | PUR | - |
| 39 | (1, 3, 0, 0, 1) | CHB | CEU,TSI,FIN | - | - | GIH |
| 40 | (1, 2, 2, 0, 0) | CHB | CEU,TSI | YRI,LWK | - | - |
| 41 | (1, 2, 1, 1, 0) | CHB | CEU,TSI | YRI | PUR | - |
| 42 | (1, 2, 1, 0, 1) | CHB | CEU,TSI | YRI | - | GIH |
| 43 | (1, 2, 0, 2, 0) | CHB | CEU,TSI | - | PUR,ACB | - |
| 44 | (1, 2, 0, 1, 1) | CHB | CEU,TSI | - | PUR | GIH |
| 45 | (1, 2, 0, 0, 2) | CHB | CEU,TSI | - | - | GIH,PJL |
| 46 | (1, 1, 3, 0, 0) | CHB | CEU | YRI,LWK,GWD | - | - |
| 47 | (1, 1, 2, 1, 0) | CHB | CEU | YRI,LWK | PUR | - |
| 48 | (1, 1, 2, 0, 1) | CHB | CEU | YRI,LWK | - | GIH |
| 49 | (1, 1, 1, 2, 0) | CHB | CEU | YRI | PUR,ACB | - |
| 50 | (1, 1, 1, 1, 1) | CHB | CEU | YRI | PUR | GIH |
| 51 | (1, 1, 1, 0, 2) | CHB | CEU | YRI | - | GIH,PJL |
| 52 | (1, 1, 0, 3, 0) | CHB | CEU | - | PUR,ACB,CLM | - |
| 53 | (1, 1, 0, 2, 1) | CHB | CEU | - | PUR,ACB | GIH |
| 54 | (1, 1, 0, 1, 2) | CHB | CEU | - | PUR | GIH,PJL |
| 55 | (1, 1, 0, 0, 3) | CHB | CEU | - | - | GIH,PJL,BEB |
| 56 | (1, 0, 4, 0, 0) | CHB | - | YRI,LWK,GWD,MSL | - | - |
| 57 | (1, 0, 3, 1, 0) | CHB | - | YRI,LWK,GWD | PUR | - |
| 58 | (1, 0, 3, 0, 1) | CHB | - | YRI,LWK,GWD | - | GIH |
| 59 | (1, 0, 2, 2, 0) | CHB | - | YRI,LWK | PUR,ACB | - |
| 60 | (1, 0, 2, 1, 1) | CHB | - | YRI,LWK | PUR | GIH |
| 61 | (1, 0, 2, 0, 2) | CHB | - | YRI,LWK | - | GIH,PJL |
| 62 | (1, 0, 1, 3, 0) | CHB | - | YRI | PUR,ACB,CLM | - |
| 63 | (1, 0, 1, 2, 1) | CHB | - | YRI | PUR,ACB | GIH |
| 64 | (1, 0, 1, 1, 2) | CHB | - | YRI | PUR | GIH,PJL |
| 65 | (1, 0, 1, 0, 3) | CHB | - | YRI | - | GIH,PJL,BEB |
| 66 | (1, 0, 0, 4, 0) | CHB | - | - | PUR,ACB,CLM,PEL | - |
| 67 | (1, 0, 0, 3, 1) | CHB | - | - | PUR,ACB,CLM | GIH |
| 68 | (1, 0, 0, 2, 2) | CHB | - | - | PUR,ACB | GIH,PJL |
| 69 | (1, 0, 0, 1, 3) | CHB | - | - | PUR | GIH,PJL,BEB |
| 70 | (1, 0, 0, 0, 4) | CHB | - | - | - | GIH,PJL,BEB,STU |
| 71 | (0, 5, 0, 0, 0) | - | CEU,TSI,FIN,GBR,IBS | - | - | - |
| 72 | (0, 4, 1, 0, 0) | - | CEU,TSI,FIN,GBR | YRI | - | - |
| 73 | (0, 4, 0, 1, 0) | - | CEU,TSI,FIN,GBR | - | PUR | - |
| 74 | (0, 4, 0, 0, 1) | - | CEU,TSI,FIN,GBR | - | - | GIH |
| 75 | (0, 3, 2, 0, 0) | - | CEU,TSI,FIN | YRI,LWK | - | - |
| 76 | (0, 3, 1, 1, 0) | - | CEU,TSI,FIN | YRI | PUR | - |
| 77 | (0, 3, 1, 0, 1) | - | CEU,TSI,FIN | YRI | - | GIH |
| 78 | (0, 3, 0, 2, 0) | - | CEU,TSI,FIN | - | PUR,ACB | - |
| 79 | (0, 3, 0, 1, 1) | - | CEU,TSI,FIN | - | PUR | GIH |
| 80 | (0, 3, 0, 0, 2) | - | CEU,TSI,FIN | - | - | GIH,PJL |
| 81 | (0, 2, 3, 0, 0) | - | CEU,TSI | YRI,LWK,GWD | - | - |
| 82 | (0, 2, 2, 1, 0) | - | CEU,TSI | YRI,LWK | PUR | - |
| 83 | (0, 2, 2, 0, 1) | - | CEU,TSI | YRI,LWK | - | GIH |
| 84 | (0, 2, 1, 2, 0) | - | CEU,TSI | YRI | PUR,ACB | - |
| 85 | (0, 2, 1, 1, 1) | - | CEU,TSI | YRI | PUR | GIH |
| 86 | (0, 2, 1, 0, 2) | - | CEU,TSI | YRI | - | GIH,PJL |
| 87 | (0, 2, 0, 3, 0) | - | CEU,TSI | - | PUR,ACB,CLM | - |
| 88 | (0, 2, 0, 2, 1) | - | CEU,TSI | - | PUR,ACB | GIH |
| 89 | (0, 2, 0, 1, 2) | - | CEU,TSI | - | PUR | GIH,PJL |
| 90 | (0, 2, 0, 0, 3) | - | CEU,TSI | - | - | GIH,PJL,BEB |
| 91 | (0, 1, 4, 0, 0) | - | CEU | YRI,LWK,GWD,MSL | - | - |
| 92 | (0, 1, 3, 1, 0) | - | CEU | YRI,LWK,GWD | PUR | - |
| 93 | (0, 1, 3, 0, 1) | - | CEU | YRI,LWK,GWD | - | GIH |
| 94 | (0, 1, 2, 2, 0) | - | CEU | YRI,LWK | PUR,ACB | - |
| 95 | (0, 1, 2, 1, 1) | - | CEU | YRI,LWK | PUR | GIH |
| 96 | (0, 1, 2, 0, 2) | - | CEU | YRI,LWK | - | GIH,PJL |
| 97 | (0, 1, 1, 3, 0) | - | CEU | YRI | PUR,ACB,CLM | - |
| 98 | (0, 1, 1, 2, 1) | - | CEU | YRI | PUR,ACB | GIH |
| 99 | (0, 1, 1, 1, 2) | - | CEU | YRI | PUR | GIH,PJL |
| 100 | (0, 1, 1, 0, 3) | - | CEU | YRI | - | GIH,PJL,BEB |
| 101 | (0, 1, 0, 4, 0) | - | CEU | - | PUR,ACB,CLM,PEL | - |
| 102 | (0, 1, 0, 3, 1) | - | CEU | - | PUR,ACB,CLM | GIH |
| 103 | (0, 1, 0, 2, 2) | - | CEU | - | PUR,ACB | GIH,PJL |
| 104 | (0, 1, 0, 1, 3) | - | CEU | - | PUR | GIH,PJL,BEB |
| 105 | (0, 1, 0, 0, 4) | - | CEU | - | - | GIH,PJL,BEB,STU |
| 106 | (0, 0, 5, 0, 0) | - | - | YRI,LWK,GWD,MSL,ESN | - | - |
| 107 | (0, 0, 4, 1, 0) | - | - | YRI,LWK,GWD,MSL | PUR | - |
| 108 | (0, 0, 4, 0, 1) | - | - | YRI,LWK,GWD,MSL | - | GIH |
| 109 | (0, 0, 3, 2, 0) | - | - | YRI,LWK,GWD | PUR,ACB | - |
| 110 | (0, 0, 3, 1, 1) | - | - | YRI,LWK,GWD | PUR | GIH |
| 111 | (0, 0, 3, 0, 2) | - | - | YRI,LWK,GWD | - | GIH,PJL |
| 112 | (0, 0, 2, 3, 0) | - | - | YRI,LWK | PUR,ACB,CLM | - |
| 113 | (0, 0, 2, 2, 1) | - | - | YRI,LWK | PUR,ACB | GIH |
| 114 | (0, 0, 2, 1, 2) | - | - | YRI,LWK | PUR | GIH,PJL |
| 115 | (0, 0, 2, 0, 3) | - | - | YRI,LWK | - | GIH,PJL,BEB |
| 116 | (0, 0, 1, 4, 0) | - | - | YRI | PUR,ACB,CLM,PEL | - |
| 117 | (0, 0, 1, 3, 1) | - | - | YRI | PUR,ACB,CLM | GIH |
| 118 | (0, 0, 1, 2, 2) | - | - | YRI | PUR,ACB | GIH,PJL |
| 119 | (0, 0, 1, 1, 3) | - | - | YRI | PUR | GIH,PJL,BEB |
| 120 | (0, 0, 1, 0, 4) | - | - | YRI | - | GIH,PJL,BEB,STU |
| 121 | (0, 0, 0, 5, 0) | - | - | - | PUR,ACB,CLM,PEL,MXL | - |
| 122 | (0, 0, 0, 4, 1) | - | - | - | PUR,ACB,CLM,PEL | GIH |
| 123 | (0, 0, 0, 3, 2) | - | - | - | PUR,ACB,CLM | GIH,PJL |
| 124 | (0, 0, 0, 2, 3) | - | - | - | PUR,ACB | GIH,PJL,BEB |
| 125 | (0, 0, 0, 1, 4) | - | - | - | PUR | GIH,PJL,BEB,STU |
| 126 | (0, 0, 0, 0, 5) | - | - | - | - | GIH,PJL,BEB,STU,ITU |

RP: Reference panel. Pop.: population. 5 groups were refer to vector *i_1_ to i_5_*, the combinations were the positive integer solutions of function “*i_1_+i_2_+i_3_+i_4_+i_5_=5*”. The populations in same one group was in sequence, for example, if *i_1_*=1 for EAS group, then the CHB was included in final reference panel, if *i_1_*=2, then CHB and CHS were included. Each combined reference panels contained 640 haplotypes.

***Supplementary Table 3: 1000G reference panels imputation performance in Han Chinese with different conditions***

| panel | MafBin | SNPNum | EmpR^2^>=0.8 | EmpR^2^<0.8 | Mean EmpR^2^ | Median EmpR^2^ | EmpR^2^ SD |
| --- | --- | --- | --- | --- | --- | --- | --- |
| 1000G panel without singleton and performed in Michigan server | 0 | 8 | 0 | 8 | 0 | 0 | 0 |
|  | 0-0.025% | 942 | 189 | 753 | 0.24964 | 2.00E-05 | 0.39211 |
|  | 0.025-0.1% | 12361 | 2510 | 9851 | 0.42574 | 0.34976 | 0.33129 |
|  | 0.1-0.5% | 26210 | 10919 | 15291 | 0.6606 | 0.72636 | 0.29101 |
|  | 0.5-1% | 8008 | 5871 | 2137 | 0.84033 | 0.93399 | 0.21504 |
|  | 1-5% | 42132 | 36956 | 5176 | 0.91814 | 0.97604 | 0.14175 |
|  | 5-100% | 426747 | 393203 | 33544 | 0.94064 | 0.98194 | 0.10939 |
| 1000G panel without singleton and performed in local server | 0 | 8 | 0 | 8 | 0 | 0 | 0 |
|  | 0-0.025% | 942 | 189 | 753 | 0.24965 | 2.00E-05 | 0.39212 |
|  | 0.025-0.1% | 12362 | 2513 | 9849 | 0.42577 | 0.34983 | 0.33129 |
|  | 0.1-0.5% | 26209 | 10920 | 15289 | 0.6606 | 0.72631 | 0.29101 |
|  | 0.5-1% | 8008 | 5871 | 2137 | 0.84033 | 0.93399 | 0.21505 |
|  | 1-5% | 42132 | 36958 | 5174 | 0.91814 | 0.97602 | 0.14175 |
|  | 5-100% | 426747 | 393214 | 33533 | 0.94064 | 0.98194 | 0.10939 |
| 1000G panel with singleton and performed in local server | 0 | 9 | 0 | 9 | 0 | 0 | 0 |
|  | 0-0.025% | 932 | 186 | 746 | 0.24789 | 2.00E-05 | 0.39216 |
|  | 0.025-0.1% | 12373 | 2514 | 9859 | 0.42509 | 0.34896 | 0.3313 |
|  | 0.1-0.5% | 26212 | 10922 | 15290 | 0.66008 | 0.72474 | 0.2913 |
|  | 0.5-1% | 8001 | 5868 | 2133 | 0.84064 | 0.93371 | 0.21448 |
|  | 1-5% | 42143 | 36928 | 5215 | 0.91766 | 0.97595 | 0.14278 |
|  | 5-100% | 426740 | 392962 | 33778 | 0.94032 | 0.9819 | 0.11007 |

EmpR^2^ means empirical-R^2^, which was used to measure the imputation accuracy using the variants that were genotyped by the Illumina Human610-Quad (610K) BeadChip.. All variants were divided into 7 MAF bins for the detailed comparison between three strategies.
